# A Multi-Scale Ecological Approach to Assessing Antimicrobial Resistance in a Freshwater Fish

**DOI:** 10.64898/2026.05.07.723562

**Authors:** John Berini, Chloe A. Fouilloux, Eric Neeno-Eckwall, Heather Alexander, Emma Choi, Grace Vaziri, Jennifer McClure, Gwen Casey, Amy Chen, Shira Dubin, Cate Patterson, Amanda K. Hund, Daniel I. Bolnick, Jessica L. Hite

**Affiliations:** Biology Department, Winona State University, Minnesota, USA; Biology Department, Carleton College, Minnesota, USA; Department of Pathobiological Sciences, University of Wisconsin-Madison, Wisconsin USA; United States Department of Agriculture, Madison, Wisconsin, USA; Department of Ecology & Evolutionary Biology, University of Connecticut, USA

**Keywords:** Antimicrobial resistance genes, environmental reservoirs, land use, deforestation, aquaculture, water quality, wild fish

## Abstract

Antimicrobial resistance (AMR) genes are increasingly recognized as an emerging environmental contaminant. Yet, the ecological mechanisms shaping their distribution across natural landscapes remain poorly understood. Here, we quantified AMR gene abundances in microbial communities sampled from wild fish from eight freshwater lakes on Vancouver Island and paired these gene-level measurements with fine-scale limnological and land-use data. Using droplet digital PCR, field surveys, and an iterative spatial forecasting framework that integrates Random Forest models with regression kriging, we explored how watershed-scale processes relate to variation in AMR genes across lakes. Our analyses reveal potential associations between elevated AMR gene levels, changes in water quality, deforestation, and geographic proximity to salmon aquaculture. By integrating data across biological and spatial scales, from genes within microbial communities to lake-level conditions and landscape patterns, this study illustrates the value of combining quantitative molecular measurements with geospatial modeling to identify environmental factors that may promote antimicrobial resistance in natural systems. Our approach provides a proof-of-concept and a general predictive framework for generating hypotheses and informing future monitoring efforts aimed at understanding, managing, and forecasting environmental reservoirs of resistance.

**Significance:** Antimicrobial resistance (AMR) genes are ancient components of environmental microbiomes. Yet, the mechanisms that generate modern hotspots of resistance across natural landscapes remain unclear. Here, we reveal how watershed-scale environmental change, including water quality metrics linked with deforestation and proximity to salmon aquaculture, predicts elevated AMR gene levels in the microbiomes of wild fish populations. By combining quantitative droplet digital PCR with ecological data and geospatial modeling, we move beyond isolated surveillance data to identify ecological mechanisms that promote antimicrobial resistance in freshwater ecosystems. This integrative approach provides mechanistic insight into why certain habitats, and the organisms within them, become reservoirs of resistance while others do not. Our findings highlight the importance of ecological context in understanding resistance evolution and offer a predictive tool for informing proactive monitoring and management strategies.

## Introduction

Drug-resistant microbes are now documented in hedgehogs^1^, bats^2^, a wide array of non-human primates^3^, and migratory birds^4–7^, far from areas with any direct antibiotic use. The mere presence of the genes that confer antimicrobial-resistance (AMR) in the microbiomes of wildlife is not inherently alarming; many of these genes are ancient, widespread components of environmental microbiomes^1,8^. The emerging concern lies in the degree to which human activities amplify, mobilize, and redistribute these genes across ecosystems, increasing opportunities for transmission among species, habitats, and biogeographic regions^9–12^. At the same time, natural systems themselves may act as long-term reservoirs, periodically introducing novel resistance variants into livestock and clinical settings^1,3,13^.

Discoveries of AMR genes in novel or unexpected hosts and environments often prompt dramatic headlines predicting the collapse of antibiotic efficacy and a return to the pre-antibiotic era. If such warnings hold even partially true, then understanding, and ultimately interrupting, the processes that spread resistance genes across environmental reservoirs is a critical global priority^1,13–15^. Such insights could advance the understanding of the ecological feedbacks that modulate bacterial evolution while also revealing practical opportunities for mitigating and preventing the spread of antimicrobial resistance. Yet, despite considerable research effort, we still lack the integrated data and computational frameworks needed to mechanistically explain why some hosts and habitats consistently function as AMR “hot spots,” while others do not^16–19^.

Traditional approaches, grounded in reductionist laboratory experiments and sparse clinical or field observations, offer crucial yet partial insights into the mechanisms driving the spread of AMR genes across natural environments^9,20–22^. While genome sequencing has revolutionized bacterial surveillance, the capacity to interpret these changes and translate genomic data into predictive models and actionable mitigation strategies remains limited^23^. Integrating genomic data with important environmental and ecological meta-data could help provide a systems-level understanding of the mechanisms driving the emergence and persistence of AMR genes in natural environments^24–26^.

As a first step toward disentangling these complex interactions, we apply an integrative framework to evaluate resistance dynamics in the gill microbiomes of wild fish across freshwater lakes on Vancouver Island. We combine microbial community profiling, long-read sequencing, targeted AMR gene quantification, and a suite of statistical tools and geospatial modeling. We focus on the three-spined stickleback (*Gasterosteus aculeatus*) as a case study. This well-studied model organism inhabits marine, brackish and freshwater habitats throughout coastal temperate regions of the Northern hemisphere^27,28^. These habitats vary in their proximity to anthropogenic activities and land-use changes (Fig. 1). For example, our study lakes on Vancouver Island range from remote, difficult to access sites to locations increasingly impacted by pronounced land use changes^29,30^.

**Figure 1.**
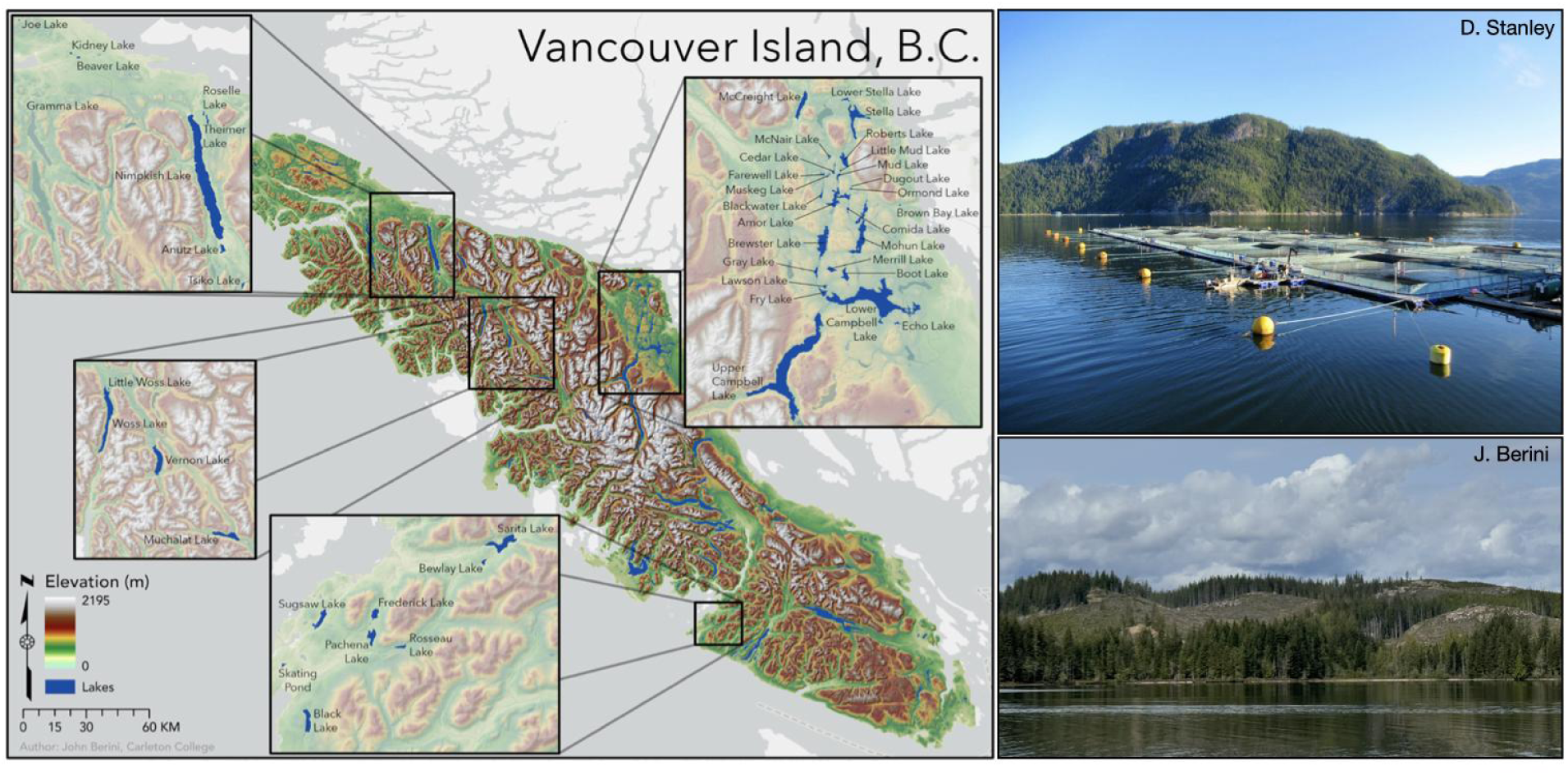
Overview of study lakes and examples of land use on Vancouver Island, British Columbia. Map of the 39 freshwater lakes sampled across multiple watersheds. Top right: Example of commercial salmon aquaculture on Vancouver Island, one of the most important salmon-producing areas on the Pacific coast (Photo credit: David Stanley, CC-BY-2.0). Bottom right: Hillslope logging above Beavertail Lake on Vancouver Island, British Columbia. Clear-cut patches are evident on the upper slopes immediately surrounding the lake, creating a sharp contrast with the intact forest cover along the shoreline and lower elevations (Photo credit: John Berini).

This region of Vancouver Island supports extensive commercial, recreational, and Indigenous salmon aquaculture making it one of the most important salmon-producing areas on the Pacific coast^31–33^. In aquaculture, antibiotic use in open-net pen systems is not spatially restricted to farms: uneaten medicated feed and antibiotic residues excreted in feces and urine can disperse into surrounding marine and freshwater habitats^34–37^. These inputs can drive local antibiotic accumulation and promote the spread of drug-resistant bacteria over distances exceeding one kilometer, depending on currents^34^. Thus, aquaculture sites can act as sources of antibiotic pressure on natural environments. Alternatively, they may also be recipients, with AMR genes from wild populations entering managed systems. Understanding this bidirectional exchange is essential for effective mitigation.

An elegant study by Larsen and colleagues illustrates this point: European hedgehogs harbored particular lineages of methicillin-resistant *Staphylococcus aureus* (MRSA) long before the antibiotic era^1^. These lineages circulated within hedgehog populations and eventually spilled over into livestock and humans. Hence, while the mere presence of AMR genes in environmental reservoirs is not inherently cause for alarm, the central challenge is determining whether, how, and when AMR genes in these reservoirs influence resistance dynamics in agricultural or clinical settings^13,38,39^.

A second land use factor that may facilitate the spread of AMR genes on Vancouver Island is logging, including widespread clear-cutting of old-growth forests^40,41^ (Fig. 1). These anthropogenic activities can reshape natural watersheds and degrade water quality through increased nutrients and sedimentation^42^. In turn, changes in water quality metrics such as chlorophyll *a* (a proxy for phytoplankton biomass, which typically increases with nutrient inputs^43,44^), dissolved oxygen, water temperature, and pH can mediate generalized stress responses in bacteria, including the upregulation of AMR genes.

AMR genes are a critical component of bacterial stress physiology, as many resistance mechanisms, including efflux pumps and membrane-associated defenses, evolved to help bacteria survive and compete with other microbes under variable and stressful environmental conditions^45–47^. For instance, both pH and oxidative stress can upregulate efflux systems^48,49^, including those associated with resistance to florfenicol and oxytetracycline. Together, the gene-environment feedbacks help explain why AMR genes are linked with eutrophic conditions in both the field and laboratory^2,50^. Although these processes are difficult to disentangle using field data alone, they highlight how local environmental conditions can interact with microbial physiology to structure the abundance and diversity of AMR genes.

Because lakes and rivers on Vancouver Island are connected by tributary networks (Fig. 1), land use changes in surrounding habitats can alter how water, nutrients, and organisms, including microbes, move across the landscape. Such connectivity can facilitate the movement of bacteria and their genes among aquatic systems^51^, providing a natural context for uncovering the ecological and evolutionary mechanisms that shape how antimicrobial resistance emerges, persists, and spreads within complex microbial communities.

## Results

### Wild fish microbiomes harbor AMR genes with relevance to aquaculture and livestock

We collected gill swab samples from a total of 195 fish across 39 lakes (Fig. 2). To characterize the overall gill microbiome, we used 16S rRNA sequencing. Next, to evaluate the antimicrobial resistome in these samples, we used long-read nanopore sequencing on a subset consisting of nine samples from three lakes, pooled among six barcodes. Using BLAST, we surveyed the CARD database to identify appropriate AMR genes for further investigation. Among the candidates were *floR*, *tap*, *mexQ/mdsB* homologs (multi-drug efflux transporters), *muxB* (a multi-drug efflux pump), *triC* (a triclosan-specific efflux protein), and *efpA* (a mycobacterial transporter). In this study, we focus on the *floR* and *tap* genes because they are particularly relevant for aquaculture. The *floR* and *tap* genes are Major Facilitator Superfamily (MFS) efflux transporters, with *floR* being specific for florfenicol, and closely related to *cml*, a chloramphenicol transporter. The *tap* gene, on the other hand, is specific for tetracyclines. Oxytetracycline, a broad-spectrum tetracycline-class drug, is widely used to treat pneumonia and bacterial enteritis (e.g., *E. coli*) in cattle, sheep, and pigs, while florfenicol is frequently used to treat respiratory diseases, particularly in swine and poultry^52^. Critically, both florfenicol and oxytetracycline are among the most widely used antibiotics in salmon aquaculture globally^53,54^ and across Canada^55^.

**Figure 2.**
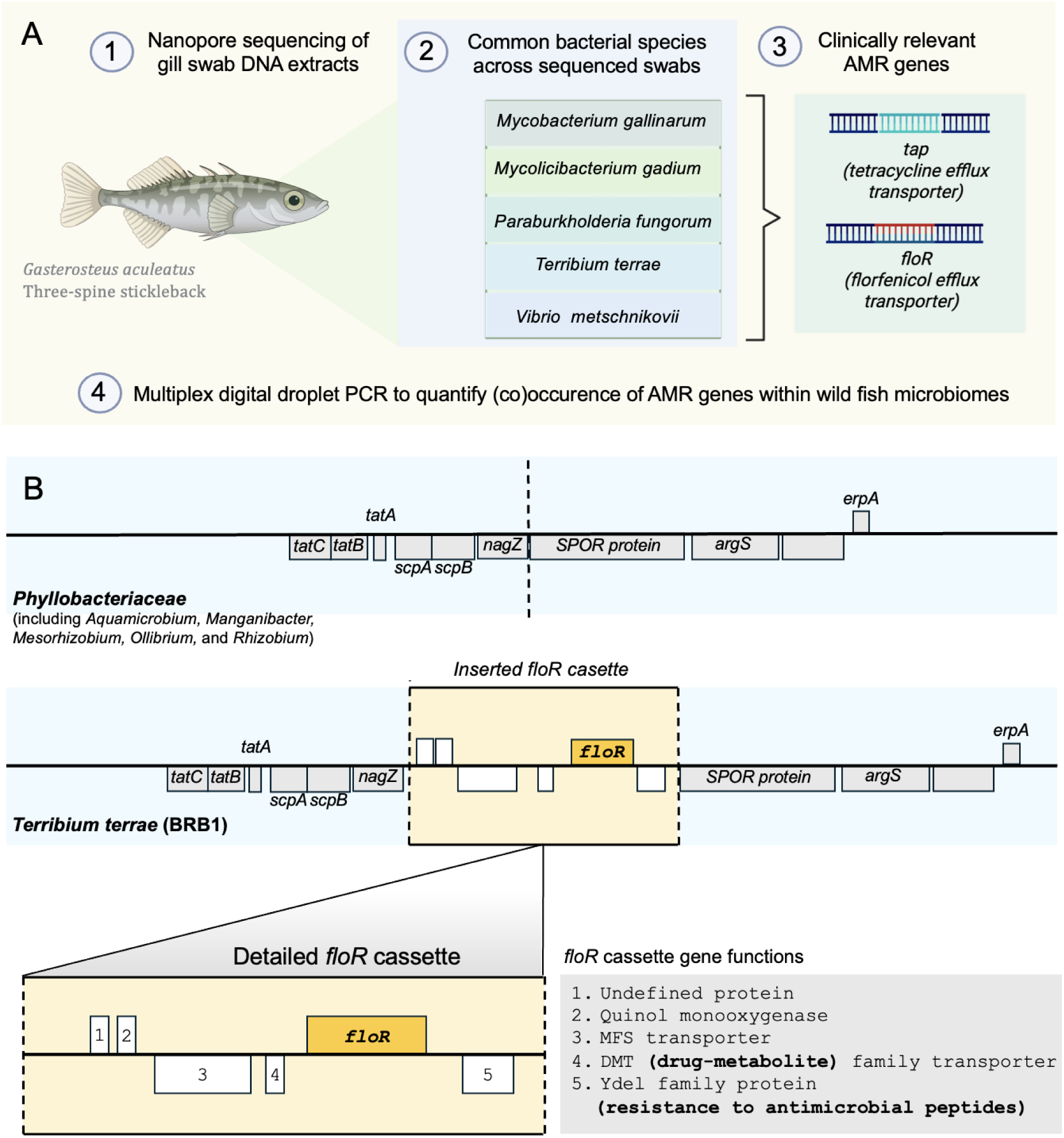
(A) Overview of the pipeline used to identify and quantify antimicrobial resistance (AMR) genes. **(1)** We extracted DNA from gill swabs collected from natural populations of three-spined stickleback (*Gasterosteus aculeatus*) in freshwater lakes across Vancouver Island. Using nanopore long-read sequencing, we identified the suite of AMR genes present in a subset of the samples. **(2)** We used Kraken to assign bacterial taxonomy and applied BLAST searches against the Comprehensive Antibiotic Resistance Database (CARD) to identify AMR genes within the sequence data. **(3)** To move beyond documenting AMR gene presence, we quantified select AMR genes that are most relevant to aquaculture, livestock, and public health with multi-plexed droplet digital PCR (ddPCR) across samples collected from multiple lakes. **Note:** Bacterial taxa and AMR genes presented here represent a subset of those identified from our samples. (**B**) **Putative bacterial hosts were inferred using a combination of sequence homology, gene–taxon co-occurrence patterns, and genomic context.** Sequence alignments indicated that the *floR*_BC_ variant most closely matched gene sequences from Gram-negative soil bacteria, including *Terribium terrae* (formerly *Mesorhizobium*), whereas the tap_BC_ gene aligned most closely with a Gram-positive actinomycete, a non-tuberculous mycobacterium in the genus *Mycolicibacterium*.

To move beyond documenting the presence of these genes, we quantified their abundance using multi-plexed ddPCR (n = 194 individual samples). To account for differences in fish biomass, which can vary depending on ecological and environmental conditions in lakes, we present size-corrected data. We distinguish the Vancouver Island, British Columbia variants identified in this study with a ‘BC’ subscript to facilitate comparisons to other published versions. Across all samples that were positive for AMR genes (total AMR concentration > 1.5 ng/uL; n = 26), the *floR_BC_* gene was more abundant (size-corrected mean 5.64 ± μL ^-1^host^-1^g^-1^, Fig. 3) than the *tap_BC_* gene (size-corrected mean 4.02 ± 0.762 SE, gene copies 0.546 SE, gene copies μL ^-1^host^-1^g^-1^ Fig. 3). These results confirm the occurrence of AMR genes conferring resistance to antibiotics commonly used to treat a wide array of bacterial infections in human and veterinary medicine, including aquaculture.

**Figure 3.**
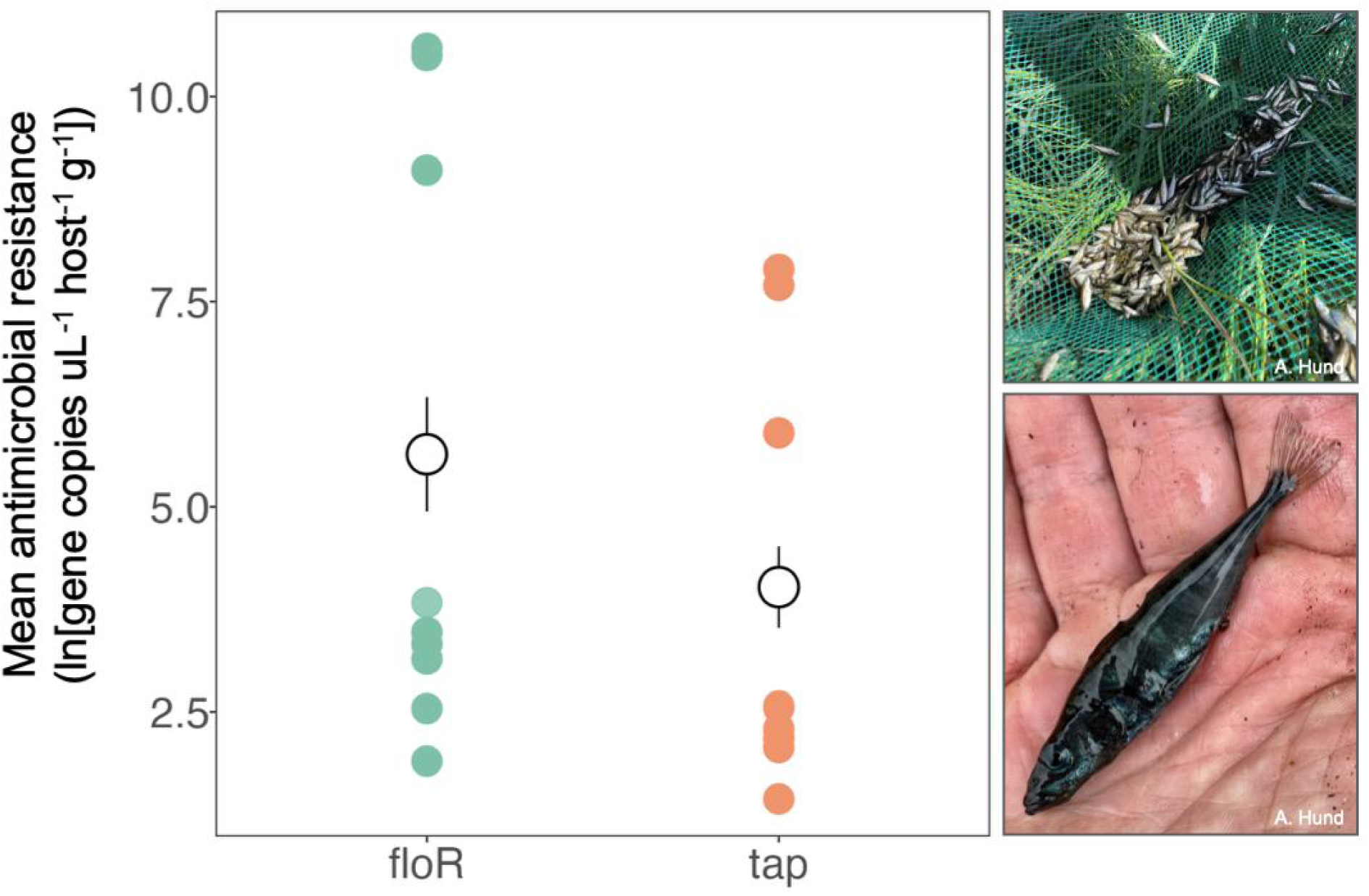
Abundance of two AMR genes conferring resistance to antibiotics commonly used in salmon aquaculture. We used multiplexed droplet digital PCR (ddPCR) to quantify the abundance of two focal genes isolated from the gill microbiomes of wild-caught three-spined stickleback fish, pictured here (n = 195) from 39 lakes across Vancouver Island. Across all AMR-positive samples (1.5 ug/nl n_Female_= 10, n_Male_= 10, 13%), the *floR_BC_* gene was more abundant (size-corrected mean 5.64 ± 1.4 SE, gene copies μL ^-1^host^-1^g^-1^) than the *tap_BC_* gene (size-corrected mean 4.02 ± 0.98 SE, gene copies μL ^-1^host^-1^g^-1^). Both genes encode efflux pumps conferring resistance to florfenicol (a phenicol-class antibiotic) and oxytetracycline (a broad-spectrum tetracycline-class drug). Both are among the most widely used therapeutants in global salmon aquaculture, including in those near our field survey lakes on Vancouver Island, British Columbia. Photo Credit: Amanda Hund.

The version of *floR* (*floR_BC_*) that we identified from the raw nanopore sequence reads shares 57% amino acid identity (74% similarity) to *pp-flo* from *Photobacterium damselae*^56^ (formerly *Pasteurella piscicida*) and 100% identity to a sequence in *Terribium terrae* KCTC 72278 (formerly: *Mesorhizobium*^57^ and identified as such in the Silva database). The sequence we identified as *tap* (*tap_BC_*) is 67% identical (82% similar) to the version in *Mycobacterium tuberculosis* with the closest match in GenBank being 98% identical to *Mycobacterium gallinarum* (Supplementary Table 1).

To explore the bacterial hosts and genome context associated with these focal AMR genes, we assembled the nanopore reads from a sample that was derived from three fish from a single lake (Brown Bay). This resulted in a 99% complete *Terribium terrae* genome consisting of two contigs and a plasmid (5,974,950 bp total) (Supplementary Fig. 1). We also retrieved over 700 kilobases of assembled mycobacterial DNA on 35 contigs whose taxonomic assignment is more difficult to ascertain. Upon inspection, the *floR_BC_* and *tap_BC_*genes both appear to be chromosomal in origin (Fig. 2B and Supplementary Fig. 2).

We identified eleven sequenced genomes in GenBank that contain a *floR* gene with 100% protein sequence identity to *floR_BC_*. Using Average Nucleotide Identity (ANI) analysis, which provides a measure of genomic similarity at the nucleotide level between genomes, we found that they are all over 99% identical to each other and to the genome we recovered from our nanopore sequencing, suggesting that they are all strains of *Terribium terrae* despite most being assigned to the genus *Mesorhizobium* in NCBI (Supplementary Fig. 3).

### Female hosts harbor the highest levels of AMR genes

Due to sex-based differences in mating behaviors, we hypothesized that males would have greater exposure to soil bacteria and thus carry higher levels of AMR genes. During the typical breeding season on Vancouver Island (June-July)^58^, males spend prolonged periods building and guarding their nests where females lay eggs^59^ (Fig. 4A). These small, tunnel-shaped structures are composed of plant debris, fine sediment, and “spiggin,” a proteinaceous glue secreted from the kidneys^60^. After the eggs are laid, males spend 3-10 days sticking their heads in these nests and fanning the eggs, which likely stirs up more sediment. These breeding behaviors could result in males having closer contact with benthic substrates and suspended sediments, potentially increasing their exposure to soil-linked environmental microbes, such as those identified in our samples (Fig. 4B).

**Figure 4.**
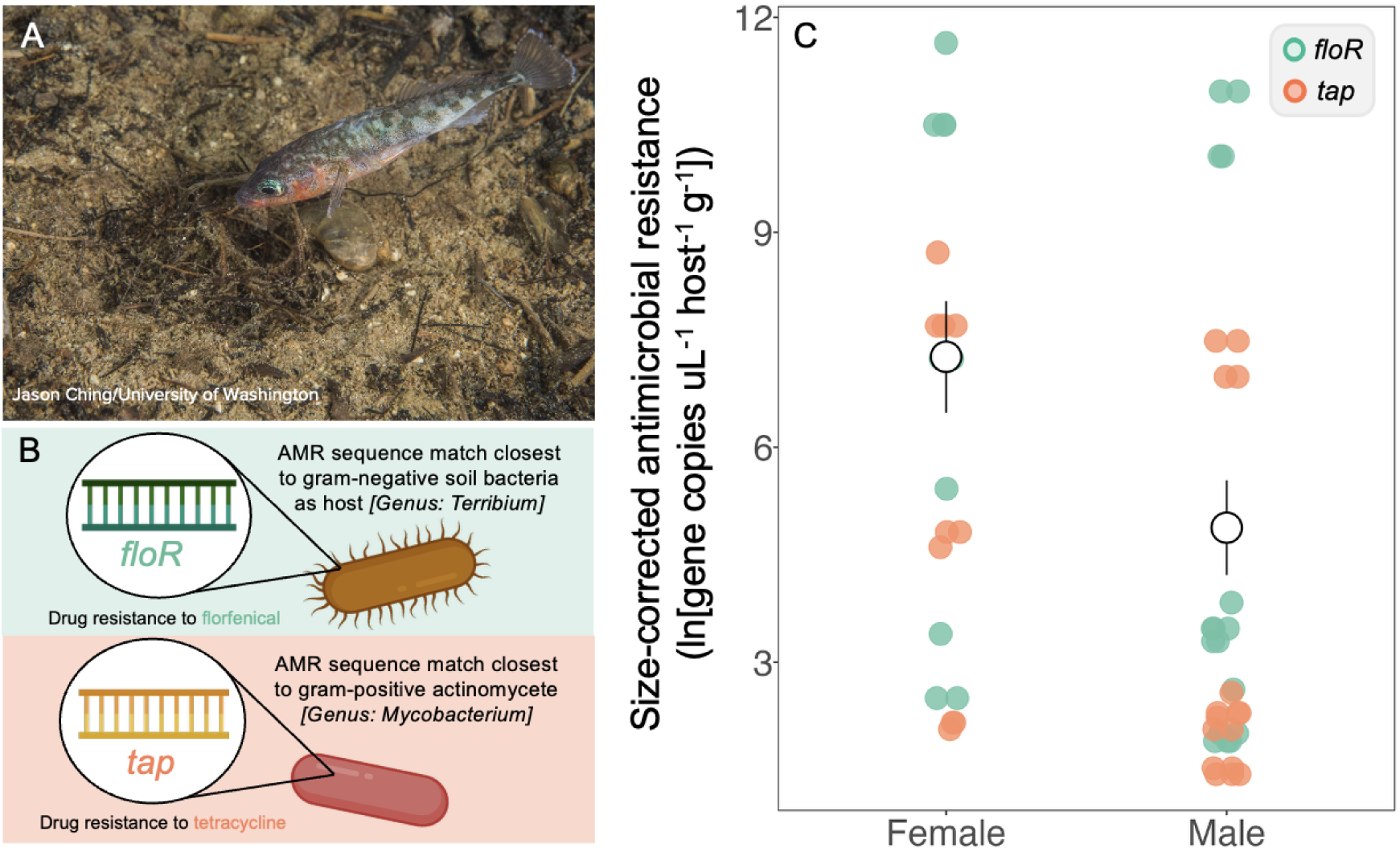
Female hosts harbor higher levels of antimicrobial resistance (AMR) genes. **(A)** Male three-spined stickleback construct nests (using a mixture of plant debris, “spiggin”, fine sediment) that are repeatedly visited by females for spawning. Photo credit: Jason Ching. (**B**) Alignment-based analyses indicated that the *floR*_BC_ variant most closely matched gene sequences from Gram-negative soil bacteria, including *Terribium terrae* (formerly *Mesorhizobium*), whereas the tap_BC_ gene aligned most closely with a Gram-positive actinomycete, a non-tuberculous mycobacterium in the genus *Mycolicibacterium*. **(C)** Across the subset of wild-caught stickleback that tested positive for AMR (n = 26), females carried significantly higher levels of AMR genes in their gill microbiomes (7.26 ± 0.806 SE) relative to males (4.88 ± 0.548 SE), even after accounting for sex-based differences in body mass (Wilcoxon rank-sum test (two-sided), W = 111, p-val = 0.047).

Contrary to our hypothesis, however, the highest levels of AMR genes were found in females. Across all lakes, female fish carried significantly higher levels of AMR genes relative to males, even after accounting for sex-based differences in body mass (Wilcoxon rank-sum test, W = 111, p-val = 0.047; size-corrected mean females 7.26 ± 0.806 SE vs. size-corrected mean males 4.88 ± 0.548 SE gene copies μL ^-1^host^-1^g^-1^, Fig. 4C).

### Land-use change linked to hot spots of AMR genes

We hypothesized that land-use intensification (aquaculture and logging) would be associated with elevated levels of antimicrobial resistance genes through at least two key pathways. We tested this idea using a geostatistical modeling framework that integrated our multiplexed ddPCR data (n = 194 swabs from individual fish, including samples that were both positive and negative for AMR genes). We compiled spatial environmental covariates compiled from publicly available databases (iMapBC, GeoBC Freshwater Atlas^61^, and NASA EarthData^62^, the ClimateNAr^63,64^ and the *lakemorpho*^65^ packages in R). We applied a three-stage modelling workflow, briefly described below (see Appendix for extended methods).

First, we applied a random forest–based preprocessing step to reduce redundancy among the large set of candidate land use, climate, and environmental covariates while retaining predictors that were consistently informative across repeated stochastic model fits. Next, using this set of refined environmental covariates, we fit and selected a parsimonious multiple regression model and then generated spatial predictions using regression kriging, which combines regression predictions with kriging of residual spatial structure not explained by covariates. Finally, we evaluated the predictive performance of the full regression–kriging workflow^66^.^67,68^ using leave-one-out cross-validation (LOOCV)^69–71^.

The most parsimonious model explaining spatial variation in AMR gene levels across our study region included four covariates: distance to old-growth forests, distance to the nearest lake, mean number of frost-free days in September, and tertiary species cover (F*_4,23_* = 5.7790, adjusted R² = 0.4145 p = 0.0023; Table 1). Residual variation was low (σ = 1.05), indicating that the selected predictors explained most of the large-scale environmental variation in AMR across Vancouver Island. The combined regression–kriging surface revealed strong spatial heterogeneity in AMR genes across the island (Fig. 5). Predicted levels of AMR genes were highest in coastal, low-elevation regions, particularly along the eastern coastline and near areas of aquaculture activity, whereas interior and high-elevation watersheds exhibited lower levels of predicted AMR genes.

**Figure 5.**
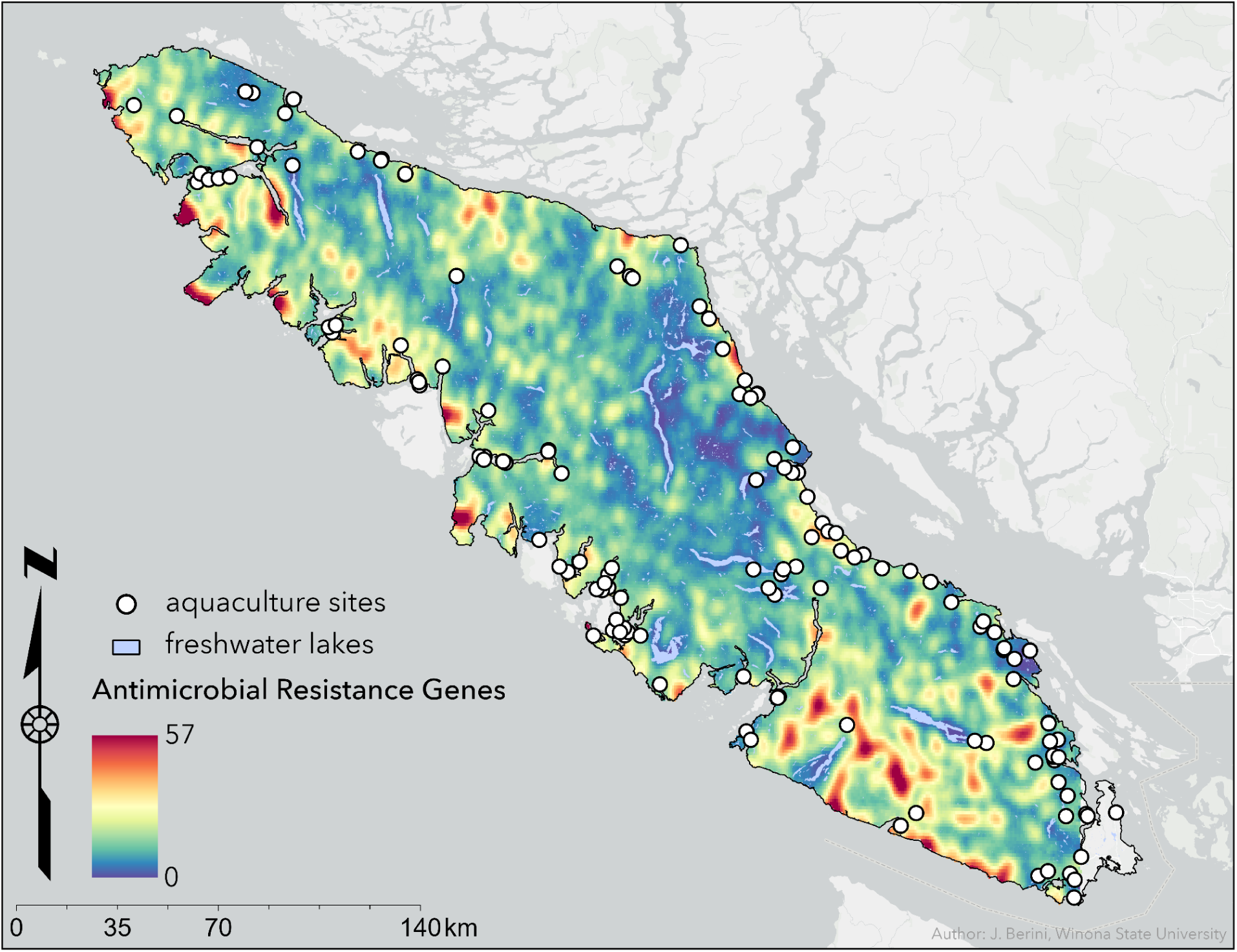
Spatial variation in antimicrobial resistance (AMR) genes. To understand potential links between land use and the pronounced spatial variation in the abundance of AMR genes in natural populations of three-spined stickleback across Vancouver Island, British Columbia, we combined our genomic data with spatial environmental covariates using a mixed-effects regression kriging approach. These geospatial models incorporate watershed-scale land use patterns, including metrics of forest alteration (distance to ancient forest and tertiary tree species cover) and geographic proximity to commercial salmon aquaculture farms, to predict AMR gene distributions. Predicted levels of AMR genes are highest (red) in southwestern and southern portions of Vancouver Island, with elevated concentrations (orange–yellow) extending along portions of the western and southwestern coasts. Isolated hotspots of elevated AMR genes are also evident along the southeastern coast. In contrast, interior and northern mountainous areas, as well as much of the northeastern coast, exhibit consistently lower predicted values (blue). While aquaculture sites (white circles) are distributed around the island’s perimeter, the spatial alignment of AMR hotspots with metrics of forest loss and landscape alteration suggests that broad-scale anthropogenic ecosystem degradation is an important driver of AMR gene distributions in the microbiomes of wild fish populations.

**Table 1.** Linear mixed model characterizing spatial variation in antimicrobial resistance (AMR) genes. Standardized coefficients (β), standard errors (SE), t-statistics, and p-values from a linear mixed model fitted with REML using Satterthwaite’s method for hypothesis tests of linear mixed effects models. The model was selected following comparison with a null model (random intercept only) using likelihood ratio test (χ² = 9.557, p = 0.049). Fixed effects include distance to nearest lake, tertiary species cover, distance to aquaculture, and distance to old growth forest (> 400 years). All predictors were scaled prior to model fitting. Random intercepts account for variation among bedrock geology groups (n = 8) across a sample size of 28 lakes.

| Fixed Effect | $\beta$ | SE | t | p |
| --- | --- | --- | --- | --- |
| Intercept | 0.848 | 0.284 | 2.983 | 0.089 |
| Distance to next closest lake | 0.551 | 0.257 | 2.147 | 0.050 |
| Tertiary Species Cover | 0.611 | 0.297 | 2.057 | 0.053 |
| Distance to Aquaculture | -0.029 | 0.253 | -0.117 | 0.908 |
| Distance to Ancient Forest | -0.337 | 0.292 | -1.154 | 0.260 |

To validate the model and evaluate its predictive performance relative to our dataset, we used spatial leave-one-out cross-validation (LOOCV). We found that predicted levels of AMR genes closely aligned with observed values across our study sites (Pearson’s r = 0.84, p < 0.001). Prediction error was low (RMSE = 0.77), and no systematic bias was evident in the residual distribution or in the relationship between residuals and predicted values. Spatial mapping of cross-validation residuals showed little regional clustering, indicating that the model effectively captured both environmental and spatial drivers of AMR variation across Vancouver Island (Supplementary Fig. 6).

Together, our data-guided geospatial model identified potential links between elevated AMR levels and watershed-scale land use patterns, including geographic proximity to commercial salmon farms, and logging operations, including access roads. Our approach illustrates how this general framework can be applied to other data sets to predict potential AMR hot spots and relevant environmental drivers across broader spatial scales.

### Water quality is linked with variation in the abundance of antimicrobial resistance (AMR) genes

We hypothesized that water-quality conditions could help explain the observed links between land-use intensification and AMR gene levels. To assess this possibility, we quantified key water-quality variables across a subset of the nine lakes that were positive for AMR genes. Unfortunately, water quality data for one lake was lost from the sonde interface, leaving a final sample size of eight lakes.

We focused on four water-quality metrics that are also known to induce generalized stress responses in bacteria, including the expression of AMR genes: resource availability, oxygen, temperature, and pH^17,72–75^. Each of these environmental variables can have non-linear and sometimes opposing effects on microbial physiology and gene expression. As a result, the direction and magnitude of any relationships between water quality and AMR genes were difficult to predict *a priori*, underscoring the need for an empirical, exploratory framework such as the one presented here. We collected these water quality metrics using an EXO2 multi-probe sonde (YSI Incorporated, Ohio, USA) to measure temperature-depth profiles at one meter increments to one meter above the hypolimnion. We assessed these relationships using Pearson correlations. Although we explored various path and multivariate analyses, the limited sample size (eight lakes) constrained the robustness and interpretability of these approaches.

Our data indicate relationships between the abundance of AMR genes and all four metrics of water quality (Fig. 6): resource availability (quantified here as chlorophyll *a,* a proxy for phytoplankton biomass, which typically increases with nutrient inputs; Pearson t = 1.99, r = 0.63, p = 0.093, Fig. 6A), oxygen (dissolved oxygen (DO); Pearson t = -2.07, r = −0.65, p = 0.084, Fig. 6B), water temperature (Pearson t= -2.27, r = −0.68, p = 0.064, Fig. 6C), and pH (Pearson -2.69, r = −0.74, ***p = 0.036, Fig. 6D). Given the limited sample size and the variability inherent in field data, only the pH pattern was statistically significant. Nonetheless, the surprisingly strong correlation coefficients, especially from field survey data of 8 lakes, indicate the *potential* for biologically relevant relationships.

**Figure 6.**
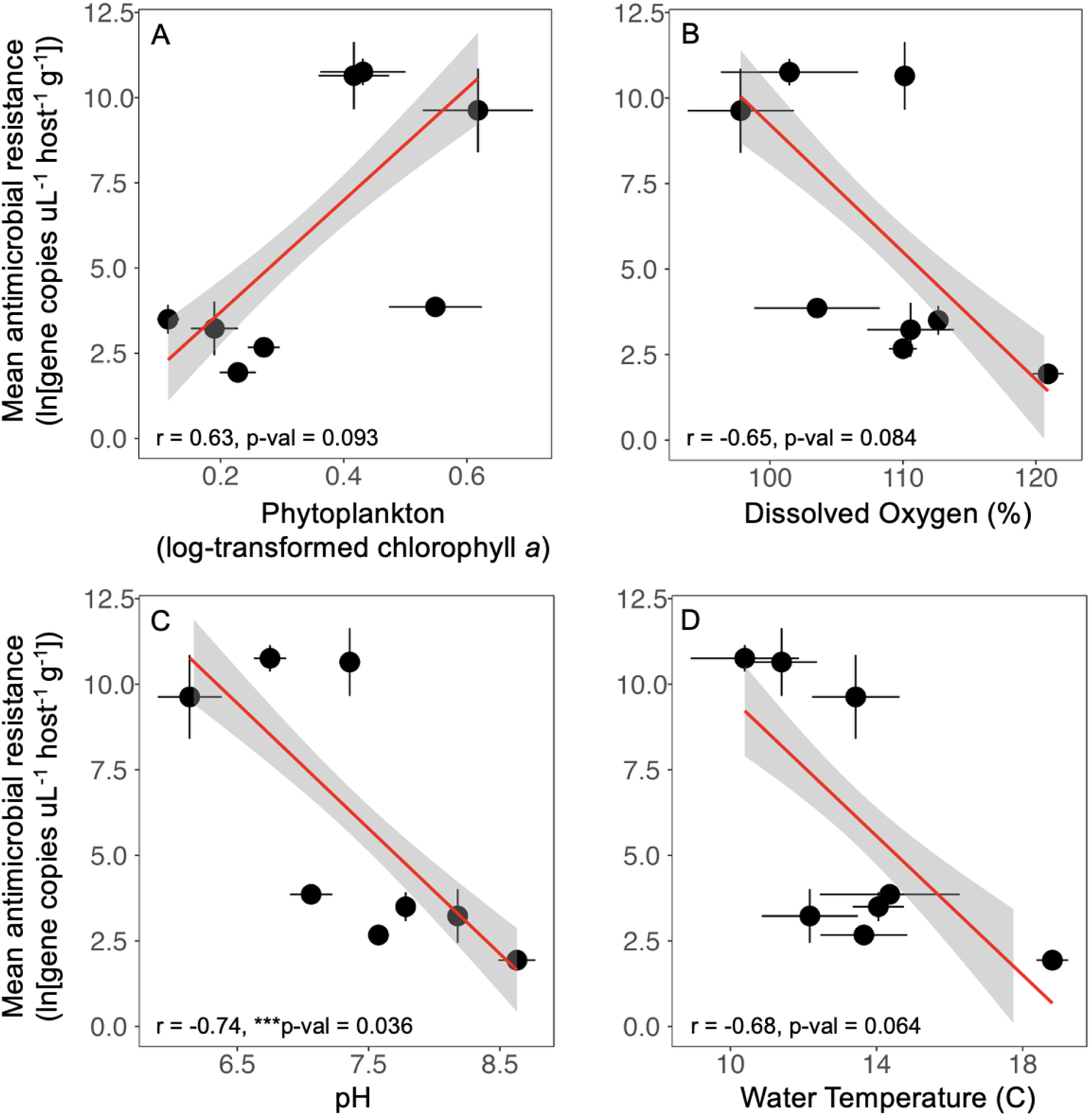
Water quality is linked with variation in the abundance of antimicrobial resistance (AMR) genes. AMR gene abundance was positively associated with multiple indicators of water quality, including (**A**) chlorophyll *a* (a proxy for phytoplankton biomass, which typically increases with nutrient inputs), (**B**) dissolved oxygen, (**C)** pH, and (**D**) temperature. Among these field-derived patterns, and small sample sizes, only the relationship with pH was statistically significant. However, the overall strength of the correlations suggests potentially meaningful ecological links with all water quality metrics and AMR genes. Data reflect the average (Mean ± SE) of individual fish sampled within each of eight lakes across Vancouver Island. Note. We limited these analyses to fish that were positive for AMR genes (total AMR concentration > 1.5 ng/uL; n = 26). Water quality measurements were obtained from integrated sonde profiles of the water column. Two-sided Pearson correlations are reported on the bottom-left corner of each plot.

These results suggest that elevated levels of AMR genes occur in lakes characterized by higher productivity, lower oxygen availability, and more acidic conditions. Each of these factors are commonly associated with lakes experiencing strong coupling between primary production and microbial respiration.

## Discussion

Our approach provides a proof-of-concept and a general predictive framework for linking quantitative measurements of antimicrobial resistance (AMR) genes to fine-scale ecological and land-use data. This integrative framework offers a much needed tool to test hypotheses and inform future monitoring of environmental reservoirs of resistance^16,24,76^. Our sample size is admittedly limited, thus these results should be interpreted cautiously. Nonetheless, the ability to link microbial genes, host traits, and watershed-scale environmental variation provides a rare opportunity to evaluate how processes operating across biological and spatial scales combine to shape antimicrobial resistance in natural ecosystems. Such cross-scale integration is uncommon in studies of environmental resistomes, which often focus on either molecular mechanisms or coarse landscape correlates such as extreme weather events or climate trends^23,38^.

We also emphasize that a central goal of this study is to highlight the value (and challenges) in moving beyond the standard approach of documenting the presence of AMR genes without the essential metadata (e.g., ecological, host, environmental) required to help identify potential mechanisms driving these emergent patterns. Such insights are crucial to extend the knowledge gained from laboratory and clinical studies often focused on single bacteria in simplified communities to complex microbial communities in which the bacteria and their genes evolve and spread^16,77^.

While aquatic habitats are increasingly recognized as key reservoirs of AMR genes, including resistance mechanisms that are of major concern for human health^35,51,78,79^, most studies have focused on contrasting system types such as lakes versus rivers, or drinking water versus sewage ^80,81^. These studies provide critical insight, but we still urgently need to disentangle how limnological characteristics of habitats (e.g., productivity, nutrient status, and water quality) relate to the environmental distribution of AMR genes^16,18,38^. Our approach highlights how even modest datasets, when coupled with quantitative field data and spatial modeling, can reveal potential ecological mechanisms that structure the distribution of AMR genes across biological scales.

Our data suggest that both host traits and the bacterial taxa present in fish microbiomes contribute to variation in AMR genes across lakes. Soil-associated bacteria and female hosts carried the highest AMR gene levels. Based on gene alignments, the bacteria that are strongly associated with the focal genes are *Terribium terrae* and *Mycolicibacterium*. The strains of *Terribium terrae* present in GenBank come from around the world and a variety of sources, including soil, sewage, human blood, microbiomes of protists and mouse gut, arabidopsis roots, and lake water. Other members of the *Phyllobacteriaceae* family include organisms isolated from nitrogen-fixing root nodules, soil, aquatic environments (freshwater and marine), and in association with mammals and other organisms^82^. Non-tuberculous mycobacteria, including *Mycolicibacterium*, are widespread in soils, sediments, and freshwater biofilms, and have been previously reported in fish gut microbiomes^83,84^, including the focal three-spined stickleback examined in this study^85,86^, making their appearance in gill microbiomes ecologically plausible.

The natural history of these microbes and their association with both aquatic and terrestrial environments could help clarify why AMR gene levels were higher in fish from lakes more heavily impacted by logging and water quality. Watershed disturbance from logging, for example, may enhance sediment and particulate runoff that transports primarily soil-associated, yet environmentally widespread bacteria and their AMR genes into lake ecosystems. This influx could, in turn, increase the probability of fish exposure through the gills. Because gill tissues continuously filter large volumes of water and intercept suspended particles and environmental bacteria, they serve as a natural interface through which microbes can be incorporated into their microbiome.

Additionally, sex-based differences in host physiology, immunity, or microbial filtering may shape how male and female fish contact, retain, or clear these environmentally widespread taxa^85–87^. Based on well-characterized behavioral differences between male and female stickleback, we hypothesized that males would have greater exposure to soil bacteria and thus, carry higher levels of AMR genes. Our field survey occurred during the breeding season on Vancouver Island, when male stickleback spend a significant amount of time building and guarding their nests, putting them in an environment with disturbed benthic substrates and potentially exposing them to soil-linked environmental microbes^87^.

Contrary to our hypothesis, the highest levels of AMR genes were found in females. While this finding could certainly reflect sampling bias, a potential biological explanation of this finding could arise from the physiological and energetic costs associated with repeated spawning events. Female stickleback can spawn one to two times per week during the reproductive season^59^, a level of reproductive investment that imposes substantial energetic demands and visibly reduces somatic condition relative to males^59,88^. Reproductive investment is known to trade off with immune function^89^ and to alter microbial colonization at mucosal surfaces^90^ across animal taxa. During energetically demanding reproductive periods, hosts may experience reduced clearance of environmental microbes and shifts in microbiome composition, which can indirectly influence the abundance and detectability of antimicrobial resistance genes carried by those communities.

Together, these ecological and sex-based pathways could act independently or in concert, with watershed-derived inputs supplying the microbial sources of AMR genes and other host traits governing variation in the microbiomes of stickleback fish^85–87^. While testing these potential explanations is beyond the scope of this current study, it raises intriguing and relevant questions that warrant further investigation.

## Applications for Management

To develop a spatially explicit predictive framework for antimicrobial resistance (AMR), we used a mixed-effects regression kriging process to forecast AMR gene distributions across Vancouver Island. Spatial predictions reveal pronounced geographic variation in AMR gene prevalence across the island, with notably elevated concentrations in coastal and southwestern regions and substantially lower values in interior and high-elevation watersheds. These spatial patterns correspond closely to gradients in key environmental drivers identified in the linear mixed model: distance to the nearest lake, tertiary species cover, distance to aquaculture facilities, and distance to ancient forest. Residual analysis indicates that the model captures broad-scale spatial structure effectively, though localized deviations suggest additional unmeasured fine-scale processes may influence AMR gene distributions in specific regions. These patterns align with gradients of land use changes, suggesting that stochastic processes alone cannot account for their emergence^35^. Combining quantitative ddPCR with geospatial models provides a robust framework to improve the ability to more accurately predict future hot spots of antimicrobial resistance and identify the ecological mechanisms that underpin these emergent patterns. This integrative and systems-based approach carries important implications for understanding bacterial evolution in complex microbial communities and more proactive approaches to prevent the spread of antimicrobial resistance genes.

Field surveys and geospatial modeling revealed that multiple anthropogenic drivers collectively influence AMR gene distributions across Vancouver Island’s freshwater ecosystems. While distance to aquaculture facilities was not individually significant, two covariates strongly associated with logging (distance to ancient forest and tertiary species cover) emerged as key predictors in the best-fitting model. Although distance to aquaculture was not statistically significant on its own, it was incorporated in the best-fitting model, which significantly outperformed the null model. This model improvement suggests that proximity to aquaculture contributes meaningfully to AMR distributions, even if its individual effect is modest. Altogether, these results implicate broad-scale anthropogenic landscape transformation as a primary mechanism linking human activity to distributions of AMR genes in natural systems.

Among the AMR genes we detected, *floR* and *tap* are particularly informative because their functions align with antibiotic classes widely deployed in salmon aquaculture globally and in British Colombia: florfenicol and oxytetracycline^33,54^. These genes both encode major facilitator superfamily (MFS) transporters, with *floR* conferring resistance to florfenicol^54,91^, and tap providing resistance to tetracyclines^92,93^. Delineating pathways of transmission^20,21^ or quantifying agricultural pressures in our study sites is beyond the scope of this study. However, reducing the movement of antimicrobial resistance genes both out of and into aquaculture remains a key concern for conservation and aquaculture management. Our study provides an initial step toward this goal by identifying ecological patterns that may point to underlying mechanisms requiring further investigation.

Salmon aquaculture in British Columbia has applied chemotherapeutants since the early 1990s, with sustained use of antibiotics including florfenicol and oxytetracycline prior to substantial reductions in reported antimicrobial use after the mid-2000s^94,95^. During the late 1990s and early 2000s, annual antimicrobial use in the region exceeded 20,000 kg^95^. Current estimates for the average antibiotic application rates for British Columbia are approximately 78 mg per kg of fish produced, with treatments occurring throughout the year^33^.

Efforts to reduce antimicrobial usage in aquaculture also benefit other livestock industries as well. Florfenicol is a cornerstone treatment for respiratory illnesses and infections in cattle, porcine, and poultry systems. Tetracyclines, including oxytetracycline, are widely used in human and veterinary medicine, and the presence of tetracycline resistance genes is widespread in environmental reservoirs^96^. The *tap* gene promotes resistance to tetracycline in *Mycobacterium tuberculosis*, the causative agent of tuberculosis^93,97^. The *tap* homolog present in our samples (*tap_BC_*) has 67 percent protein identity to the version in *M. tuberculosis* (*tap_TB_*) and 82.7 percent identity to the tap gene in *Mycolicibacterium fortuitum* (*tap_FR_*) (Supplementary Table 1). This latter variant is of interest because while *tap_TB_* is specific to tetracycline, *tap_FR_* also provides some resistance to streptomycin, gentamicin, and other antibiotics^92^.

It is possible that *tap_BC_* has altered substrate specificity as well and highlights the potential for freshwater ecosystems to act as environmental reservoirs of AMR genes with relevance to human and animal health, perhaps even harboring ones that are more effective or have a broader range of specificity than ones that have been identified in known pathogens. This concern is particularly acute for taxa such as *Mycobacterium*, in which horizontal transfer of resistance and virulence genes is a major concern for both livestock and public health^98^. Beyond antibiotic resistance, tap homologs may also function in the export of toxic metabolic by-products, with antimicrobial resistance emerging as a secondary consequence of this activity^93,99^. This dual functionality may facilitate the persistence of tap-like genes even in the absence of sustained antibiotic exposure.

Both the *floR_BC_* and *tap_BC_* genes are chromosomally encoded major facilitator superfamily (MFS) transporters. This class of transporters is present in bacteria, archaea, and eukarya, and is responsible for the transport of nutrients and metabolites across cell membranes^100–102^ . Efflux systems commonly contribute to intrinsic or baseline resistance phenotypes and can have pleiotropic roles in cellular physiology, including detoxification of metabolic by-products. Genes with such pleiotropic functions are expected to experience stronger constraints on horizontal transfer, as disruption of regulatory or genomic context may incur fitness costs^103^. Consistent with this framework, the chromosomal association of tap_BC_ and floR_BC_ corresponds with resistance emerging *de novo* through standing genetic variation or regulatory modulation rather than dissemination through mobile genetic elements.

Despite the absence of obvious mobile genetic elements in the immediate neighborhood of the *floR_BC_* gene, it seems likely that it has been integrated into the chromosome along with several other genes. In comparison to other members of the *Phyllobacteriaceae* family which lack *floR_BC_*, *Terribium terrae* contains a potential AMR gene cassette consisting of six genes, including a drug/metabolite family transporter, a potential antimicrobial peptide resistance protein, and two MFS transporters (including *floR_BC_*). This cassette is inserted between two genes (*nagZ* and a SPOR-domain protein) in a highly conserved region of these chromosomes (Fig. 2B); related strains that contain *floR* homologs appear to have them in other chromosomal locations. Similarly, the tap genes in mycobacteria appear to be located on the chromosome without obvious nearby mobile elements (Supplementary Fig. 2).

It is important to underscore that chromosomal association should not be interpreted as evidence of immobility. Although rapid dissemination of drug resistance is driven primarily by horizontal gene transfer and mobile genetic elements, many resistance determinants are chromosomally encoded and persist as part of standing genetic variation within microbial populations. Comparative genomic analyses demonstrate that chromosomal resistance genes can be transferred onto plasmids, and vice versa, through recombination and mobilization mediated by integrases, insertion sequences, and other genetic machinery^103^. Thus, the current genomic context of *floR_BC_* and *tap_BC_* reflects their state in the sampled hosts, not a fixed evolutionary endpoint. This genetic architecture informs, but does not resolve, key questions concerning which environments or hosts represent sources and which represent sinks of AMR genes.

### Linking Ecological Mechanisms with Bacteria and their Genes

Geospatial modelling linked logging to the abundance and distribution of AMR genes across Vancouver Island, with distance to ancient forest and tertiary species cover emerging as key spatial predictors. To help identify the potential ecological mechanisms driving emergent patterns in antimicrobial resistance genes, we focused on classical metrics of water quality^43,104^ that could shape the abundance of AMR genes within complex microbial communities. The expression of drug-resistance genes is an evolutionarily conserved microbial response to physiological stress^1,8^, suggesting that changes in water quality, such as shifts in resource availability, dissolved oxygen, temperature, and pH, could alter microbial stress levels^17,73,105,106^ and, in turn, influence AMR gene abundance in the microbiomes of wild stickleback fish. Because land-use change, including deforestation, can modify these limnological features, we hypothesized that lakes experiencing greater disturbance, for example due to increased allochthonous inputs from logging operations and access roads, would favor higher AMR gene levels.

Our results support this hypothesis and join other field studies demonstrating strong correlations between AMR genes and water quality conditions, dissolved oxygen, and specific nutrients. Additionally, laboratory studies indicate that oxidative and pH stress can induce or favor efflux-pump-based resistance. Oxidative stress under low dissolved oxygen can upregulate efflux pumps, including those associated with the two focal genes in this study: *floR* and *tap*. Furthermore, acidic conditions may reduce pump efficiency, potentially intensifying selective pressures for additional resistance mechanisms. Although these mechanisms remain difficult to disentangle with field data alone, the results illustrate how local environmental conditions can interact with microbial physiology to shape resistomes in freshwater ecosystems.

## Conclusions

Together, our field surveys, quantitative genomic data, and geospatial modeling help explain why some hosts and environments emerge as ‘hot spots’ of antimicrobial resistance. Elevated levels of antimicrobial resistance (AMR) genes in the gill microbiomes of wild stickleback populations were higher in female hosts, carried by two common bacteria often associated with soil microbiomes, and associated with the hallmarks of lower water quality including higher primary productivity, lower pH, and lower dissolved oxygen. These ecological factors could help explain why AMR genes are more abundant in lakes more heavily impacted by land-use pressures, including logging, access roads, and commercial salmon aquaculture. By connecting these cross-scale patterns, from genes to hosts and landscapes, this study underscores the importance of systems-based approaches to understand how bacteria and their genes emerge and spread in complex microbial communities.

## Data Availability

All data and code supporting the findings of this study will be deposited in the Environmental Data Initiative repository upon acceptance for publication. Sequence reads used in this study are available in the NCBI Sequence Read Archive under accession number PRJNA1438775. Limnological and host-specific data and associated analysis code are currently hosted at GitHub (https://github.com/jhite-eco-epi/AMR-land-use-water-quality-and-wildfish) and will be archived in a permanent repository prior to publication. Code used for geospatial modeling is available at GitHub (https://github.com/jberini/vancouver-island-amr). Geospatial datasets are available upon request.

## Methods

### Field Surveys

During the breeding season (June – July 2023), we sampled gill microbiomes from a total of 195 individual stickleback fish from 39 lakes spanning Vancouver Island’s major watersheds (Fig. 1). Collections were done with approval from the British Columbia Ministry of Lands Forest and the Environment (NA23-787881), the Huu-ayaht First Nation, and an IACUC approval from the University of Connecticut (protocol A21-025).

### Sampling stickleback gill microbiomes

We collected gill microbiome samples using sterile flocked swabs (Puritan, Cat. #25-3318-U) and preserved in RNALater to stabilize nucleic acids for downstream genomic analyses^107^. All materials were prepared in advance to minimize handling time. Prior to sampling, a 1.2-mL cryovial prefilled with 0.5 mL RNAlater (Thermo Fisher Scientific) was opened and placed on a clean surface with the cap oriented threads-up to avoid contamination.

For each fish, a sterile swab was opened immediately before use. Following sterile techniques, a handler gently restrained the fish and elevated its head above the water surface. While ensuring that the gill filaments were not touched, the operculum was carefully opened to expose the gill arches. The sterile swab was quickly inserted between the gill filaments and rotated to collect microbial material; sampling was continued until the swab tip became visibly pink, indicating successful acquisition of oxygenated epithelial cells rather than blood.

Between fish, all reusable instruments (e.g., scissors, forceps) and any potentially contaminating work surfaces were sterilized with 80% ethanol and briefly flamed to eliminate residual DNA and microbial material. This procedure followed standard aseptic practices designed to minimize cross-sample contamination in microbiome studies. Fish were euthanized immediately after swabbing.

Following sample collection, the swab was placed tip-down into the cryovial containing RNALater. Sterile scissors were used to trim the swab shaft flush with the top of the cryovial to ensure secure closure. The cryovial was then tightly sealed to prevent leakage or contamination and labeled if not pre-labeled. Any swab that contacted a non-sterile surface (e.g., the handler’s glove, fish skin, or boat surfaces) was discarded and replaced with a new sterile swab to maintain aseptic sampling conditions required for accurate bacterial community analyses. Completed samples were placed in sealed containers and kept on ice during field collection. Upon return to the laboratory, all samples were stored at −20 °C until processing.

### Microbiome Analyses

#### DNA Extraction

To preserve longer DNA fragments for nanopore sequencing, we initially chose the Zymo high molecular weight extraction kit (Zymo Research, Cat. #D6060). RNALater was removed from the swabs (n = 45) by diluting with PBS, centrifuging, and pipetting off the supernatant. 200 uL of DNA/RNA shield was added to each sample, and they were placed in a BioSpec Mini-BeadBeater-24 for 1 minute at 2400 rpm without beads. Samples were centrifuged, and the supernatant was removed to be added back after the lysozyme step. The swabs were washed with 1 ml PBS followed by the lysozyme and proteinase K treatments as described in the manufacturer’s instructions for microbial lysis (Zymo Research).

We adopted a faster extraction method (Qiagen - QiaAmp PowerFecal Pro; Cat. #51804) to process the remainder of the samples (n = 160) after the target genes were identified by Nanopore sequencing. The swab tubes were centrifuged, the RNALater was removed, and the pellet and swab was incubated for 30 minutes at room temperature in 1 mL of 200 ug/mL Proteinase K in PBS. The tubes were centrifuged, the supernatant removed, and the swabs were transferred to bead tubes; CD1 solution was used to transfer the pellet from the swab tube into the extraction tube. The samples were vortexed for 10 minutes at maximum speed, and the rest of the protocol followed the manufacturer’s instructions. DNA concentration was quantified using a Nanodrop, and the samples were normalized to 10 ng/μL.

#### 16S rRNA amplicon sequencing

High-throughput sequencing of the 16S rRNA gene was performed to characterize the gill microbiome on a subset of samples (n=45). The V4 hypervariable region was amplified using a dual-indexed primer system^108^. PCR reactions were performed in 25 μL volumes containing 19 μL AccuPrime Pfx SuperMix (Invitrogen Cat. #12344040), 1 μL each of 10 μM forward and reverse primers, 2 μL template DNA (10 ng/μL), and 2 μL DMSO. The thermocycling conditions varied from the published protocol, using 35 cycles instead of 30 to enhance detection of rare species. The PCR products were verified by 1.5% agarose gel electrophoresis, normalized using SequalPrep Normalization Plates (Invitrogen A1051001), and pooled and quantified by qPCR using the KAPA Library Quantification Kit (Roche KAPA code: KK4824) and Qubit 1x dsDNA HS Assay Kit (Illumina). The final library was diluted to 6 pM containing 10% PhiX control (Illumina FC-110-3001) and sequenced on an Illumina MiSeq platform using V2 chemistry (Illumina MS-102-2003) to generate paired-end 250 bp reads.

These FASTQ files were processed using the DADA2 plugin within QIIME2 (v2024.2.0) and the amplicons were classified against the Silva reference sequence database^109^. ASV counts corresponding to *Mesorhizobium* and *Mycobacterium* were determined and plotted against the ddPCR quantifications of the appropriate AMR genes (*floR* and *tap*, respectively) (Supplementary Fig. 4).

#### Nanopore sequencing

To identify the diversity of antimicrobial resistance genes in our samples, we used longread Nanopore sequencing with a PromethION flow cell on a Nanopore P2 Solo sequencer (Oxford Nanopore Technologies, Oxford UK). 400 ng of purified DNA was sequenced following the protocol associated with the native barcoding kit SQK-NBD114.96. We focused on a sample that consisted of pooled DNA extracts of three different fish from a single lake (Brown Bay) where the sequencing results had the greatest abundance of bacterial reads as determined by Kraken 2^110^. This approach yielded 9.9 gigabases of total sequence data from which we recovered 178 megabases of non-host DNA after aligning the reads to the *Gasterosteus aculeatus* genome (GCF_964276395.1) with minimap2 (v.2.26-r1175) and recovering the unmapped reads with samtools (v.1.18). These reads were assembled with flye (v.2.9.2-b1786), yielding 8.3 megabases of assembled DNA sequence.

To find, quantify, and categorize AMR genes, the raw reads were formatted into a BLAST database^111^ that was subsequently queried by the entries of the CARD database^112^. This analysis resulted in the identification of homologs to *floR* and *tap* within our sequenced data. These genes were subsequently localized to the assembled contigs, revealing that *floR*_BC_ is present on one of three contigs that comprise a nearly complete version of *Terribium terrae* (formerly: *Mesorhizobium*), a Gram-negative soil bacterium^57,113^, and *tap* is localized on a contig that is most similar to *Mycobacterium gallinarum*, a high-GC Gram-positive actinomycete.

#### Quantifying antimicrobial resistance (ARM) genes

After identifying the presence of these focal genes, we used ddPCR to quantify their abundance across fish populations. An alignment of *floR*_BC_ and *tap*_BC_ genes was made from sequences downloaded from GenBank corresponding to the best hits from a BLAST query using the sequences recovered from the Nanopore data. Primers were chosen based on conserved regions of these genes, with a desired amplicon in the range of 150-300 nucleotides. The probe primers were chosen to anneal to a conserved sequence in the amplicon with an annealing temperature approximately 5℃ higher than the amplification primers (Table 2).

**Table 2.**
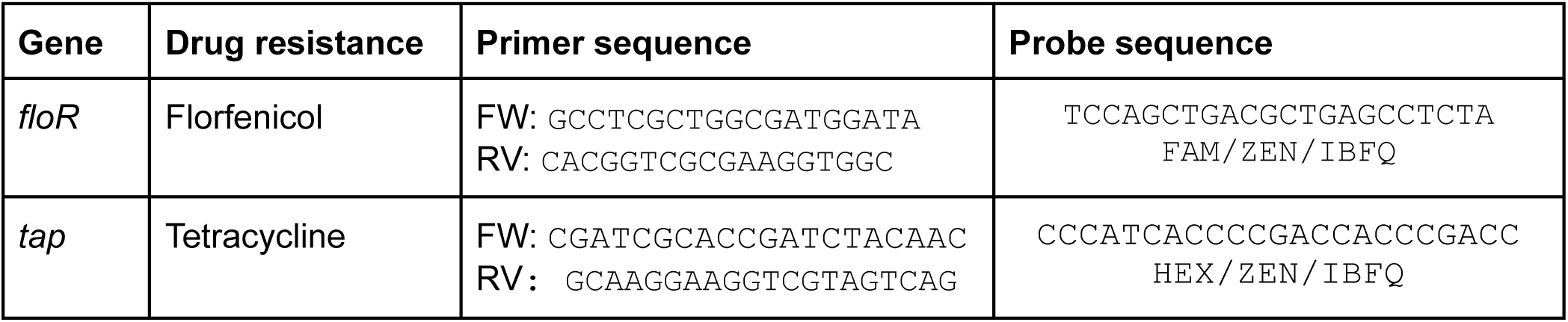
Primer and probe sequences for target AMR genes.

#### Digital Droplet PCR (ddPCR)

Before running ddPCR, amplicons were confirmed to be of the appropriate band size (*floR*_BC_ = 306bp, *tap*_BC_ = 166bp) by visualization on an agarose gel. DNA was diluted to 10ng/uL to standardize input quantity. In order to quantify AMR gene concentration in DNA extracts, we designed a multiplexed ddPCR assay where the *floR*_BC_ probe-primer was labeled with a 5’ FAM fluorophore and the *tap* primer was labeled with a 5’ HEX fluorophore. Probe-primers were ordered from Integrated DNA Technologies (IDT), these included the 5’ fluorophores specified above with a 3’ Iowa Black FQ (IBFQ) quencher and a proprietary internal ZEN quencher. ddPCR was performed using Bio-Rad’s QX-200 Droplet Digital PCR System. Each 20µl reaction was composed of 10µl of 2X ddPCR™ supermix for probes (no dUTPs, BioRad) reagent mix, FAM- and HEX-labeled probes at 250nM final concentration, forward and reverse amplicon primers at a final concentration of 900nM, and 8µl of DNA template (or RNase/DNase-free water for NTC). Droplets were generated, and 40 µl of the resulting emulsion was manually transferred to a 96-well PCR plate (Bio-Rad), which was heat-sealed with a foil cover. The droplets were then subject to thermocycling using a Bio-Rad C1000 thermocycler with a ramp rate of 1°C/s using the following specifications: a 10 min enzyme activation step at 95°C, followed by 40 cycles of 30s at 94°C (denaturation) and 1 min at 60°C (annealing/extension), followed by 10 min hold at 98°C.

Following thermocycling, the droplets were immediately read with Bio-Rad’s Droplet Reader. Absolute quantification of target gene copies (reported in concentration ng/µL) was done with default ABS settings in QuantaSoft Analysis Pro 2.0 software (Bio-Rad). Only reactions containing ≥10,000 droplets were included for further analysis^114^. As an additional quality control measure to reduce the chance of false positives, we restricted our analyses to samples above a detection threshold (total antimicrobial resistance gene concentration > 1.5 ng/uL). Based on this threshold, the final sample size for fish with positive values of AMR genes included 26 fish from eight lakes. This substantially reduced our initial sample size of 205 fish from the 47 lakes we surveyed. We acknowledge that our sample size is limited hence these results should be interpreted cautiously.

## Note

### Geostatistical Methods Overview

To characterize the drivers of antimicrobial resistance across Vancouver Island, we employed a mixed effects regression kriging framework. We first developed a deterministic trend model using linear mixed-effects models to test specific environmental and anthropogenic factors, while incorporating bedrock geology as a random effect to control for underlying bedrock porosity and potential groundwater connectivity. To account for remaining spatial dependency, we modeled the regression residuals using a spherical variogram and interpolated them across the study area via kriging. The final predictive surface was generated by integrating these kriged residuals with the regression trend, effectively merging broad-scale drivers with localized spatial patterns. We validated the robustness and stationarity of this framework using a spatial leave-one-out cross-validation (LOOCV) with fixed variogram parameters, ensuring that our findings regarding the influence of aquaculture and landscape variables remained stable across the study region.

### Geospatial Dataset

The initial set of covariates used as potential predictors of landscape-level variation in antimicrobial resistance (AMR) captured four major dimensions of spatial heterogeneity across Vancouver Island: (1) lake morphometry and geometry, (2) land cover, forest structure, and land use, (3) climate and weather, and (4) proximity to human activity and infrastructure.

Although we initially compiled 277 covariates estimated or measured across 39 lakes on Vancouver Island, we retained 19 of these covariates for subsequent covariate selection and modeling. Altogether, the retained covariates help characterize spatial variation in anthropogenic drivers, forest structure and composition, lake morphometrics, and climate.

We obtained data describing lake morphometry and geometry (1) from the GeoBC Freshwater Atlas^61^, and climate and weather variables (3) from the ClimateNA package in R^64^ . Because morphometric data were incomplete for several study lakes, we used the *lakemorpho* package in R to estimate mean and maximum depth^65^. To validate these modeled estimates, we compared them with pre-existing provincial data using Pearson correlations, confirming that they were within an acceptable range of error. We compiled variables related to land cover, forest structure, and land use (2), as well as proximity to human activity and infrastructure (4), from the iMapBC interface, except for NDVI and EVI indices, which we obtained from NASA’s EarthData platform^62^. We summarized all spatial data within a 1,000 m buffer surrounding each lake, calculating mean, minimum, and maximum values for each covariate.

### 1. Covariate screening and selection

Because our initial covariate set was large and highly correlated, we employed a hypothesis-driven screening process to identify the most parsimonious drivers of AMR. We first identified a candidate shortlist of 19 predictors based on their theoretical relevance to antibiotic resistance and landscape connectivity. To evaluate these predictors while controlling for underlying geological variance, we utilized a mixed-effects modeling framework. We fit individual Linear Mixed-Effects Models (LMM) for each candidate variable using the lme4 package, incorporating bedrock geology as a random intercept. Predictors were ranked based on p-value significance and effect size estimates. This screening identified a stable set of core predictors, specifically distance to aquaculture, distance to old-growth forest, distance to nearest lake, and species cover, which were carried forward for final model fitting and spatial prediction.

### 2. Model fitting and spatial prediction

We generated spatial predictions of AMR genes across Vancouver Island using a mixed-effects regression kriging approach, a two-step process in which a regression model captures broad-scale, covariate-driven variation and kriging of the regression residuals accounts for remaining spatial autocorrelation^66^. We first fit a global regression model incorporating the screened predictors and bedrock geology as a random effect. The significance of the fixed-effects structure was validated against a null model using a Likelihood Ratio Test. We then generated a continuous prediction of our surface from this regression model across the island’s extent and calculated residuals as the difference between observed and predicted AMR levels at sampled lakes. We characterized residual spatial dependence by fitting an empirical variogram and selecting a spherical theoretical model (Supplementary Fig. 5). These residuals were interpolated across the study area using ordinary kriging, and the resulting kriged residual surface was added to the regression trend to produce a final continuous surface representing spatial variation in AMR genes.

### 3. Model validation

To evaluate predictive performance and model stationarity, we implemented spatial leave-one-out cross-validation (LOOCV). We refit the model once for each observation in the dataset (n = 28), sequentially withholding a single lake and predicting its value from the remaining data. In each iteration, the remaining observations were used to refit the regression component and compute residuals. To ensure the stability of our spatial hypothesis and avoid overfitting to local clusters, we utilized fixed variogram parameters (nugget, sill, and range) derived from the global dataset across all iterations of the cross-validation process. The resulting prediction for each withheld site was compared with its observed value. We quantified predictive performance using root mean square error (RMSE) and the Pearson correlation coefficient (r) between observed and predicted values. Together, these metrics summarize both absolute prediction error and the consistency of spatial patterning across sites.

## Results

### Best Fitting Mixed-Effects Model of ARG Variation

Our final model identified four primary predictors of AMR gene variation: distance to the nearest lake, tertiary species cover, distance to aquaculture, and distance to ancient forest. Bedrock geology was incorporated as a random effect to account for regional geological heterogeneity. The final multivariate model demonstrated a significantly better fit to the observed data than a null model containing only the random effect of bedrock geology (*χ*^2^ = 9.557, p = 0.0486). Furthermore, the final model achieved a lower Akaike Information Criterion (AIC = 100.72) than the null model (AIC = 102.28).

### Landscape-Scale Mapping and Spatial Heterogeneity

To generate a spatially continuous surface of predicted ARG across Vancouver Island, we utilized a regression-kriging framework that combined our best-fitting mixed-effects model with the spatial structure of the model residuals. The fitted spherical variogram revealed moderate spatial dependence (nugget = 0.798, partial sill = 0.161, range = 8.857), which was used to interpolate local-scale variations not captured by the environmental covariates alone. The resulting prediction surface (Supplementary Fig. 5) reveals strong spatial heterogeneity in ARG concentrations across the study region. Predicted values are highest in the low-elevation, coastal regions, particularly along the eastern margin of the island where both human infrastructure and hydrological connectivity are densest.

### Model Validation and Predictive Performance

We assessed the reliability of our hypothesized drivers using spatial leave-one-out cross-validation (LOOCV). This validation served as a rigorous test of whether the specific environmental and geological variables identified in our hypothesis could independently predict ARG levels at unsampled locations. The model demonstrated a positive correlation between observed and predicted values (Pearson’s R = 0.22; Supplementary Fig. 6a), with an RMSE of 1.16.While the correlation coefficient reflects the inherent "noise" and complexity of AMR across the Vancouver Island landscape, the fact that a model built on a small set of specific landscape and geological predictors maintains a positive relationship in a cross-validation framework supports the validity of our hypothesized drivers.

Diagnostic analysis of the residuals further confirms this reliability - the error distribution remained consistent across the range of predicted values (Supplementary Fig. 6b) and a histogram of the residuals was well-centered at zero (Supplementary Fig. 6c), indicating no systematic flaws in the model’s logic. Furthermore, spatial mapping of the cross-validation residuals (Supplementary Fig. 6d) exhibited no regional clustering. This lack of spatial bias suggests that the model’s performance is not driven by spatial autocorrelation, but rather by the identified relationship between AMR, human influences on the local landscape, hydrology, and geology.

---

**Figure S1.**
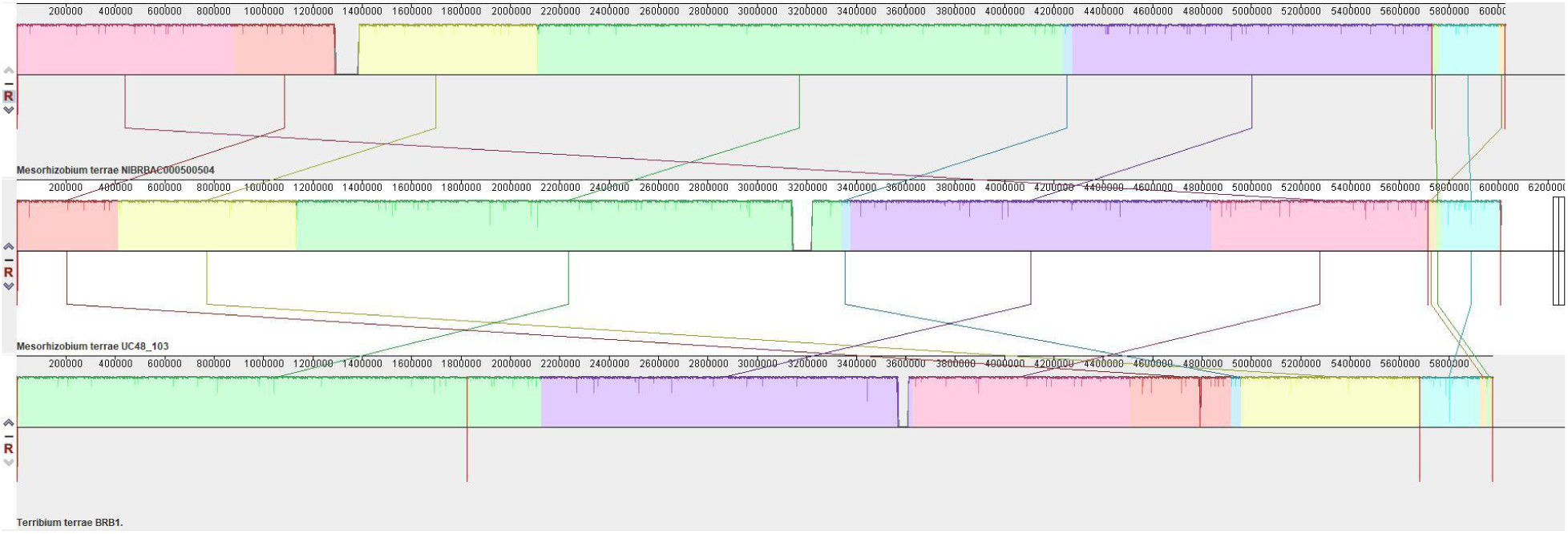
Three *Terribium terrae* genomes aligned by Mauve.|. *Terribium terrae* genomes NIBRBAC000500504 (top), UC48_103 (middle), and the metagenomic assembly BRB1 (bottom) broadly share sequence identity and context throughout the genome. Approximately 200 bp gaps are present at the ends of the BRB1 contigs relative to the other strains.

**Figure S2.**
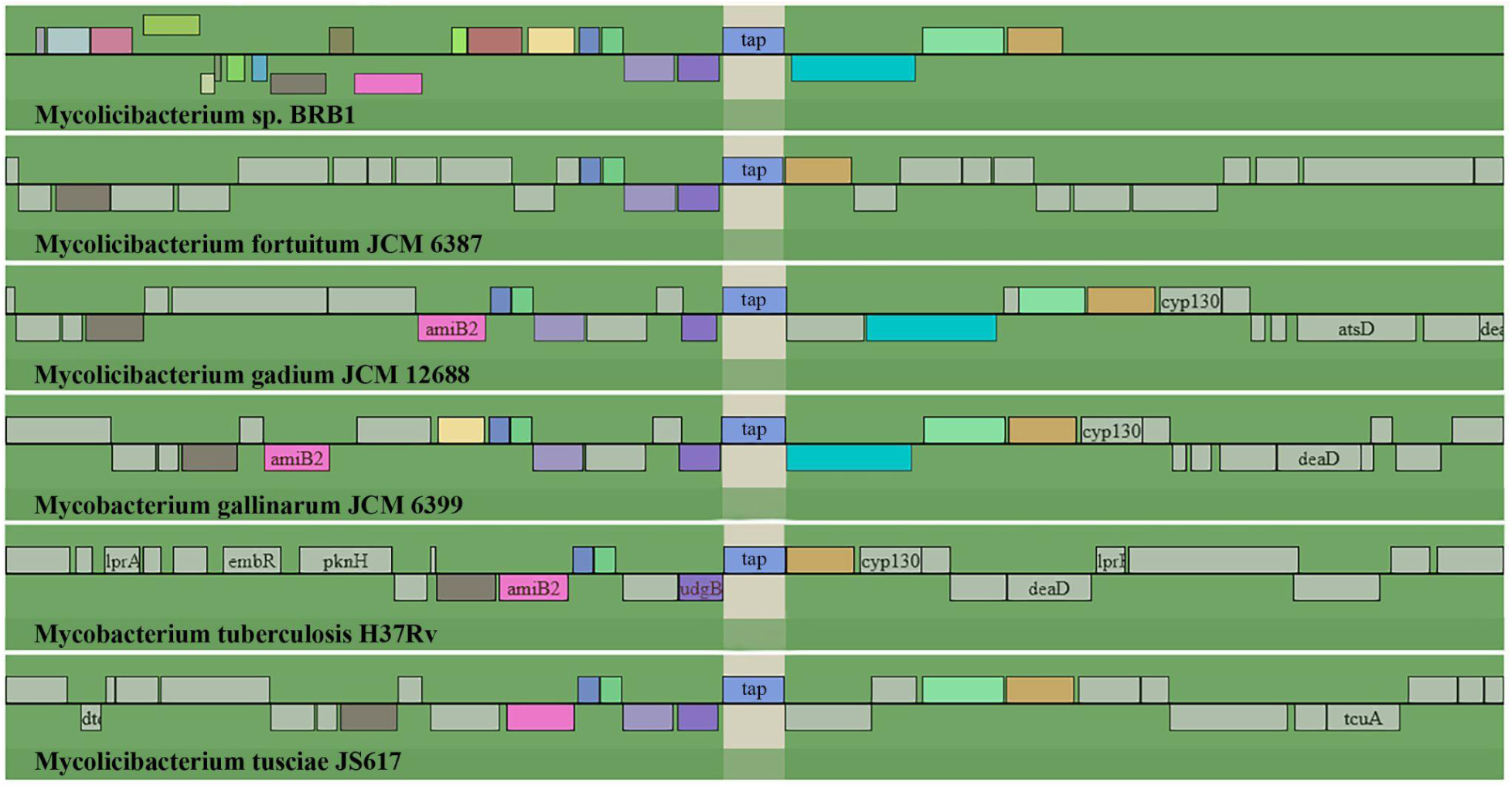
The *tap* gene is located in the same genomic region in several mycobacterial species. Colored features indicate genes that are shared in the same region as *tap_BC_*.

**Figure S3.**
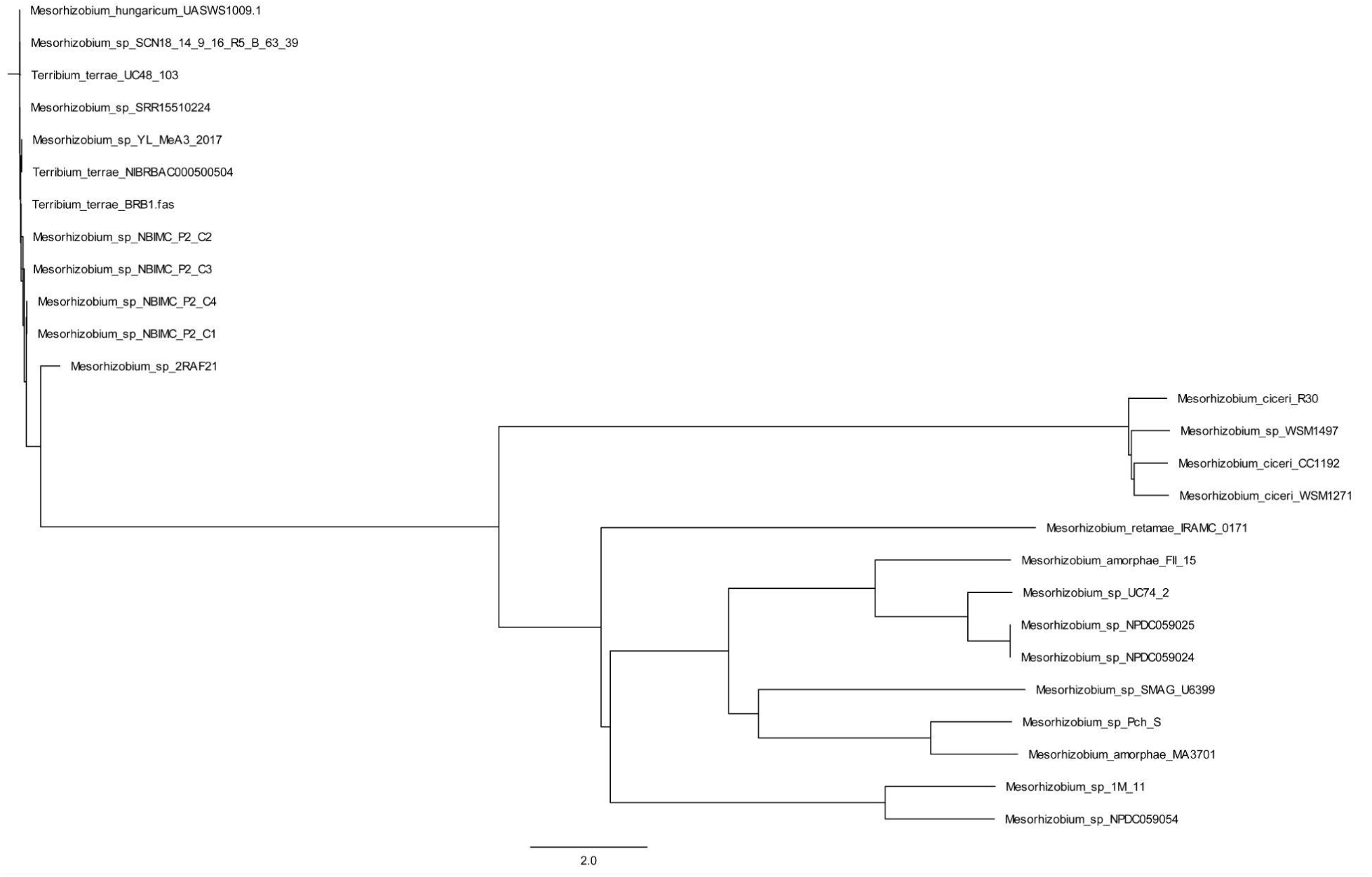
ANI analysis of floR-containing *Mesorhizobium* and *Terribium terrae* strains. Genomes from *Mesorhizobium* and *Terribium* isolates that contain homologs to the *floR* gene were downloaded from GenBank and ANI analysis was performed. All the strains containing a *floR* gene with 100% protein identity to *floR_BC_* were determined to be *Terribium terrae*, clustering within 99% identity to each other. Isolates identified as various mesorhizobia species with *floR* gene variants formed separate clusters.

**Figure S4.**
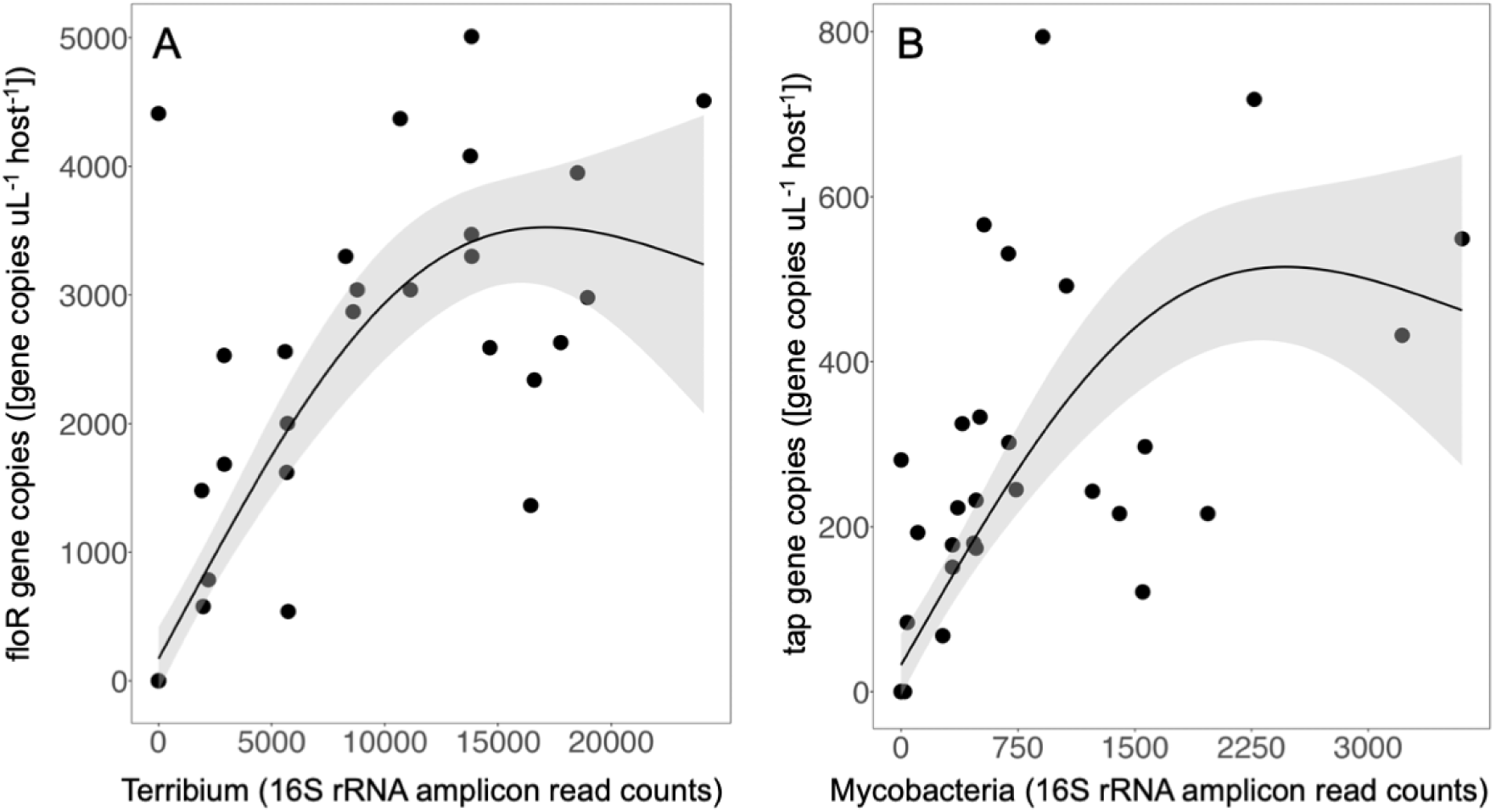
Relationship between AMR genes and putative host strains. To investigate the relationship between the amount of the AMR genes *floR* and *tap*, and the amount of the putative host strains (*Terribium* and mycobacteria, respectively), we compare the number of 16S rRNA reads that correspond to these genera with the ddPCR quantification of the AMR genes. There is a positive correlation between *Mesorhizobium* and *floR_BC_* and between mycobacteria and *tap_BC_*. The fact that these widespread bacteria are often associated with terrestrial soil microbiomes raises questions concerning potential aquatic-terrestrial linkages.

**Figure S5.**
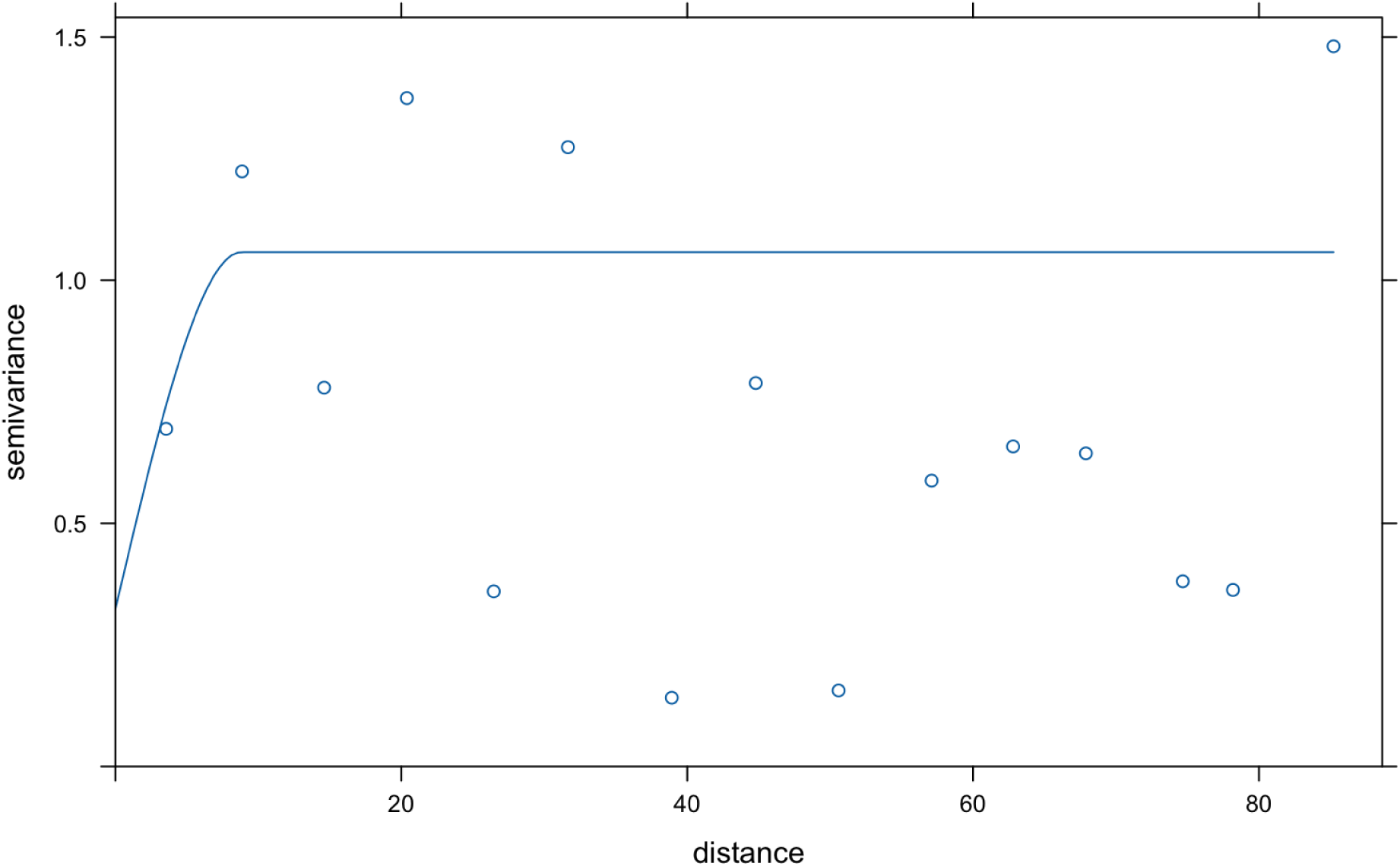
Empirical and fitted variogram of AMR model residuals. The semivariance (y-axis) is plotted against the lag distance (x-axis, in decimal degrees) to evaluate the spatial structure of AMR variation after accounting for environmental and geological fixed effects. The empirical variogram (points) was fitted with a spherical model (solid line) to determine the parameters for regression kriging. The model indicates a nugget of 0.798, representing fine-scale variation or measurement error, and a partial sill of 0.161, which describes the spatially structured component of the variance. The spatial range of 8.857 indicates the distance at which model residuals become spatially independent, suggesting that localized factors continue to influence ARG distribution beyond the broad-scale environmental covariates.

**Figure S6.**
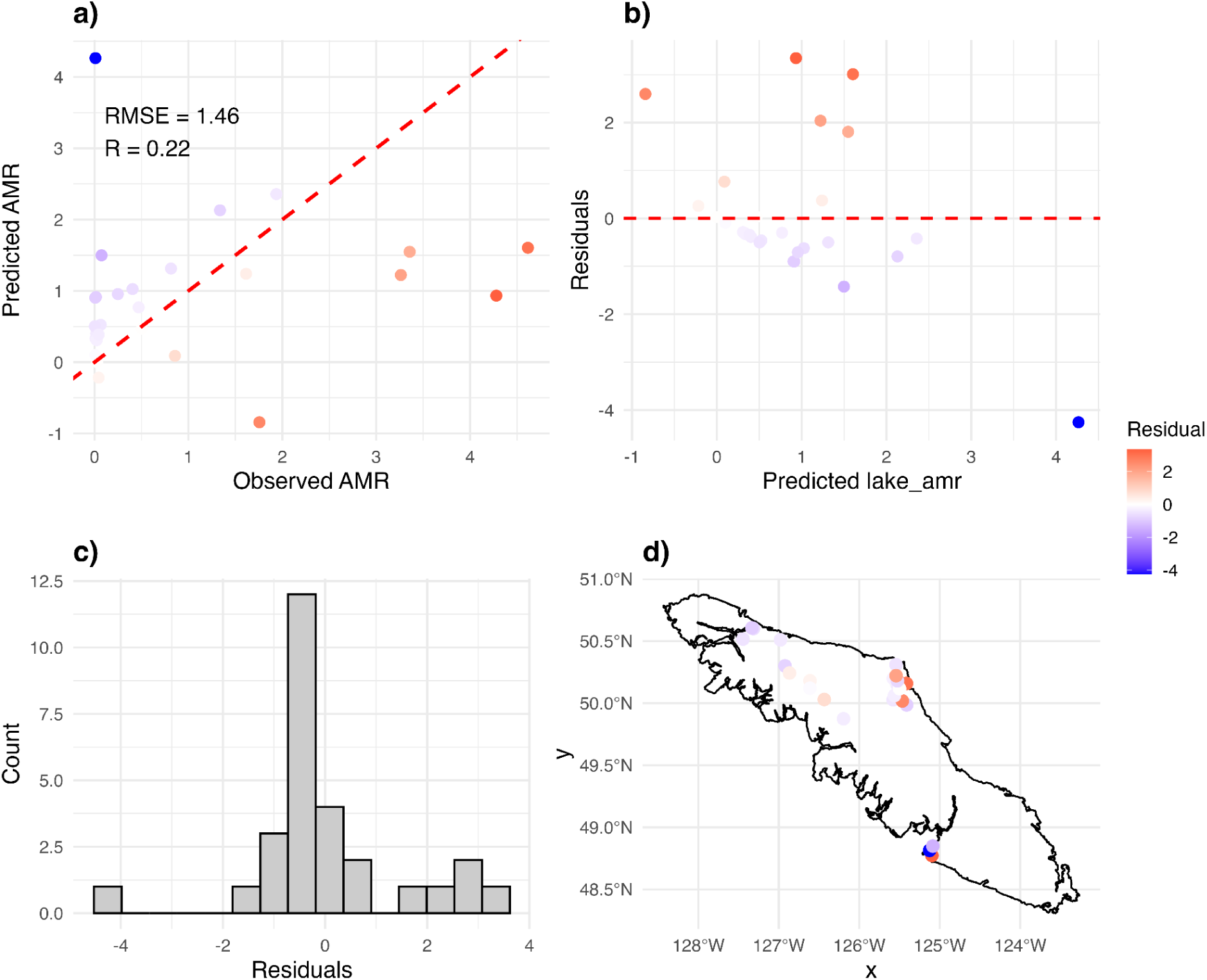
Diagnostics for spatial leave-one-out cross-validation (LOOCV) evaluating the predictive performance of the regression–kriging model across Vancouver Island (n = 39 sites). The model showed strong predictive performance (R = 0.22, RMSE = 1.46), with predicted AMR gene abundance closely tracking observed values across all sites (a). Residuals exhibited no systematic bias with respect to predicted values (b) and approximated a normal distribution (c). The spatial distribution of residuals (d) revealed minimal clustering, indicating that the regression–kriging framework effectively captured both environmental gradients and residual spatial autocorrelation in AMR gene abundance.

**Table S1:**
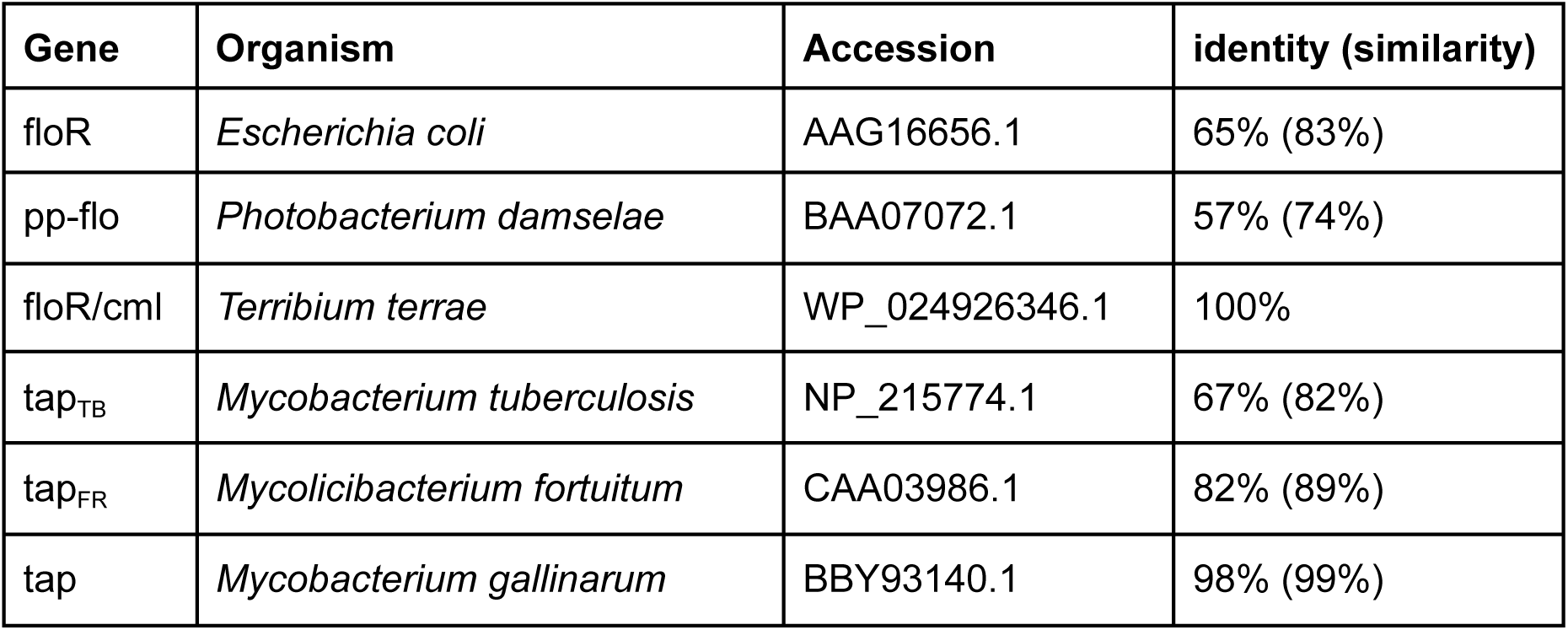
floR and tap genes from other organisms. Relationships between the versions of *floR* and *tap* identified in this study to versions reported in GenBank.

## References

1. Larsen, J. et al. Emergence of methicillin resistance predates the clinical use of antibiotics. Nature 602, 135–141 (2022).

2. Li, X., Mowlaboccus, S., Jackson, B., Cai, C. & Coombs, G. W. Antimicrobial resistance among clinically significant bacteria in wildlife: An overlooked one health concern. Int. J. Antimicrob. Agents 64, 107251 (2024).

3. Weiss, D. et al. Antibiotic-resistant Escherichia coli and class 1 integrons in humans, domestic animals, and wild primates in rural Uganda. Appl. Environ. Microbiol. 84, (2018).

4. Smith, H. G., Clarke, R. H., Larkins, J.-A., Bean, D. C. & Greenhill, A. R. Wild Australian birds and drug-resistant bacteria: characterisation of antibiotic-resistant *Escherichia coli* and *Enterococcus* spp. Emu 119, 384–390 (2019).

5. Lim, H. et al. Ecological diversity of migratory birds and their associated bacterial species in South Korea: A preliminary study including antimicrobial resistance profiles. Vet. Sci. 12, 1157 (2025).

6. Rapi, M. C. et al. Resisting the final line: Phenotypic detection of resistance to last-resort antimicrobials in Gram-negative bacteria isolated from wild birds in northern Italy. Animals (Basel) 15, 2289 (2025).

7. Sato, G., Oka, C., Asagi, M. & Ishiguro, N. Detection of conjugative R plasmids conferring chloramphenicol resistance in *Escherichia coli* isolated from domestic and feral pigeons and crows. Zentralbl. Bakteriol. Orig. A 241, 407–417 (1978).

8. D’Costa, V. M. et al. Antibiotic resistance is ancient. Nature 477, 457–461 (2011).

9. Zhu, Y.-G. et al. Microbial mass movements. Science 357, 1099–1100 (2017).

10. Storteboom, H., Arabi, M., Davis, J. G., Crimi, B. & Pruden, A. Identification of antibiotic-resistance-gene molecular signatures suitable as tracers of pristine river, urban, and agricultural sources. Environ. Sci. Technol. 44, 1947–1953 (2010).

11. Lima, T., Domingues, S. & Da Silva, G. J. Manure as a Potential Hotspot for Antibiotic Resistance Dissemination by Horizontal Gene Transfer Events. Vet. Sci. 7, (2020).

12. Knapp, C. W., Dolfing, J., Ehlert, P. A. I. & Graham, D. W. Evidence of increasing antibiotic resistance gene abundances in archived soils since 1940. Environ. Sci. Technol. 44, 580–587 (2010).

13. Woolhouse, M., Ward, M., van Bunnik, B. & Farrar, J. Antimicrobial resistance in humans, livestock and the wider environment. Philos. Trans. R. Soc. Lond. B Biol. Sci. 370, 20140083 (2015).

14. Depta, J. & Niedźwiedzka-Rystwej, P. The phenomenon of antibiotic resistance in the polar regions: An overview of the global problem. Infect. Drug Resist. 16, 1979–1995 (2023).

15. World Health Organization, Global Antimicrobial Resistance Use Surveillance System (GLASS) (2020) GLASS Whole- Genome Sequencing Surveillance Antimicrobial Resistance, World Health Organization.

16. Bottery, M. J., Pitchford, J. W. & Friman, V.-P. Ecology and evolution of antimicrobial resistance in bacterial communities. ISME J. 15, 939–948 (2021).

17. Kelbrick, M., Hesse, E. & O’ Brien, S. Cultivating antimicrobial resistance: how intensive agriculture ploughs the way for antibiotic resistance. Microbiology 169, (2023).

18. Nickodem, C. A. et al. Soil management strategies drive divergent impacts on pathogens and environmental resistomes. Sci. Rep. 15, 43215 (2025).

19. Wale, N. et al. Resource limitation prevents the emergence of drug resistance by intensifying within-host competition. Proc. Natl. Acad. Sci. U. S. A. 114, 13774–13779 (2017).

20. Lanzas, C., Davies, K., Erwin, S. & Dawson, D. On modelling environmentally transmitted pathogens. Interface Focus 10, 20190056 (2020).

21. Dawson, D., Rasmussen, D., Peng, X. & Lanzas, C. Inferring environmental transmission using phylodynamics: a case-study using simulated evolution of an enteric pathogen. J. R. Soc. Interface 18, 20210041 (2021).

22. Ghaly, T. M. & Gillings, M. R. New perspectives on mobile genetic elements: a paradigm shift for managing the antibiotic resistance crisis. Philos. Trans. R. Soc. Lond. B Biol. Sci. 377, 20200462 (2021).

23. Beltrán de Heredia, I., Alkorta, I., Garbisu, C. & Ruiz-Romera, E. A practical framework for environmental antibiotic resistance monitoring in freshwater ecosystems. Antibiotics (Basel) 14, 840 (2025).

24. Pradier, L. & Bedhomme, S. Ecology, more than antibiotics consumption, is the major predictor for the global distribution of aminoglycoside-modifying enzymes. Elife 12, (2023).

25. Bustamante, M. et al. An eco-evolutionary perspective on antimicrobial resistance in the context of One Health. iScience 28, 111534 (2025).

26. Hiltunen, T., Virta, M. & Laine, A.-L. Antibiotic resistance in the wild: an eco-evolutionary perspective. Philos. Trans. R. Soc. Lond. B Biol. Sci. 372, 20160039 (2017).

27. Reid, K., Bell, M. A. & Veeramah, K. R. Threespine stickleback: A model system for evolutionary genomics. Annu. Rev. Genomics Hum. Genet. 22, 357–383 (2021).

28. Barber, I. & Scharsack, J. P. The three-spined stickleback-*Schistocephalus solidus* system: an experimental model for investigating host-parasite interactions in fish. Parasitology 137, 411–424 (2010).

29. Parisien, M.-A. et al. Abrupt, climate-induced increase in wildfires in British Columbia since the mid-2000s. Commun. Earth Environ. 4, 309 (2023).

30. Muise, E. R. et al. Cumulative and component impacts of the human footprint on remotely sensed biodiversity indicators using dissimilarity to high integrity reference states. Int. J. Appl. Earth Obs. Geoinf. 144, 104899 (2025).

31. Atlas, W. I. et al. Indigenous systems of management for culturally and ecologically resilient pacific salmon (Oncorhynchus spp.) fisheries. Bioscience 71, 186–204 (2021).

32. Quinn, T. P. Changing themes in pacific salmon research and conservation. Rev. Fish. Sci. Aquac. 1–25 (2025).

33. Jonah, L., Hamoutene, D., Kingsbury, M., Johnson, L. & Fenton, A. J. A data compilation of antibiotic treatments in Canadian finfish aquaculture from 2016 to 2021 and the cumulative usage of antibiotics and antiparasitic drugs at marine sites. Environ. Rev. 32, 334–349 (2024).

34. Buschmann, A. H. et al. Salmon aquaculture and antimicrobial resistance in the marine environment. PLoS One 7, e42724 (2012).

35. Vittecoq, M. et al. Antimicrobial resistance in wildlife. J. Appl. Ecol. 53, 519–529 (2016).

36. Burridge, L., Weis, J. S., Cabello, F., Pizarro, J. & Bostick, K. Chemical use in salmon aquaculture: A review of current practices and possible environmental effects. Aquaculture 306, 7–23 (2010).

37. Chen, J. et al. Antibiotics and food safety in aquaculture. J. Agric. Food Chem. 68, 11908–11919 (2020).

38. Martínez, J. L. & Baquero, F. What are the missing pieces needed to stop antibiotic resistance? Microb. Biotechnol. 16, 1900–1923 (2023).

39. Sanz-García, F. et al. Translating eco-evolutionary biology into therapy to tackle antibiotic resistance. Nat. Rev. Microbiol. 21, 671–685 (2023).

40. Dhari, J. Old growth protections in British Columbia: A comparative analysis. (2023).

41. Sutherland, I. J. Long-term recovery of ecosystem services following forest harvest in coastal temperate rainforests of Vancouver Island, British Columbia, Canada. (2015).

42. León-Muñoz, J. et al. The combined impact of land use change and aquaculture on sediment and water quality in oligotrophic Lake Rupanco (North Patagonia, Chile, 40.8°S). J. Environ. Manage. 128, 283–291 (2013).

43. Filazzola, A. et al. A database of chlorophyll and water chemistry in freshwater lakes. Sci. Data 7, 310 (2020).

44. Sterner, R. W. *In situ*-measured primary production in Lake Superior. J. Great Lakes Res. 36, 139–149 (2010).

45. Darby, E. M. et al. Molecular mechanisms of antibiotic resistance revisited. Nat. Rev. Microbiol. 21, 280–295 (2023).

46. Bonilla, C. Y. Generally stressed out bacteria: Environmental stress response mechanisms in Gram-positive bacteria. Integr. Comp. Biol. 60, 126–133 (2020).

47. Dawan, J. & Ahn, J. Bacterial stress responses as potential targets in overcoming antibiotic resistance. Microorganisms 10, 1385 (2022).

48. Poole, K. Bacterial stress responses as determinants of antimicrobial resistance. J. Antimicrob. Chemother. 67, 2069–2089 (2012).

49. Nové, M., Kincses, A., Molnár, J., Amaral, L. & Spengler, G. The role of efflux pumps and environmental pH in bacterial multidrug resistance. In Vivo 34, 65–71 (2020).

50. Thevenon, F., Adatte, T., Wildi, W. & Poté, J. Antibiotic resistant bacteria/genes dissemination in lacustrine sediments highly increased following cultural eutrophication of Lake Geneva (Switzerland). Chemosphere 86, 468–476 (2012).

51. Kraemer, S. A., Barbosa da Costa, N., Oliva, A., Huot, Y. & Walsh, D. A. A resistome survey across hundreds of freshwater bacterial communities reveals the impacts of veterinary and human antibiotics use. Front. Microbiol. 13, 995418 (2022).

52. NARMS. 2019 NARMS Update: Integrated Report. (2019).

53. Lulijwa, R., Rupia, E. J. & Alfaro, A. C. Antibiotic use in aquaculture, policies and regulation, health and environmental risks: a review of the top 15 major producers. Rev. Aquac. 12, 640–663 (2020).

54. Fernández-Alarcón, C. et al. Detection of the floR gene in a diversity of florfenicol resistant Gram-negative bacilli from freshwater salmon farms in Chile. Zoonoses Public Health 57, 181–188 (2010).

55. Murphy, G. M. *Review of Antibiotic Resistance Genes (ARGs) in Salmon Aquaculture and Empirical Data on Spatial and Seasonal Trends in the Bay of Fundy*. (Fisheries and Oceans Canada, Ottawa, 2022).

56. Kim, E. & Aoki, T. Sequence analysis of the florfenicol resistance gene encoded in the transferable R-plasmid of a fish pathogen, *Pasteurella piscicida*. Microbiol. Immunol. 40, 665–669 (1996).

57. Li, Y. et al. Phylogenomic analyses and reclassification of the *Mesorhizobium* complex: proposal for 9 novel genera and reclassification of 15 species. BMC Genomics 25, 419 (2024).

58. Ishikawa, A. & Kitano, J. Diversity in reproductive seasonality in the three-spined stickleback, *Gasterosteus aculeatus*. J. Exp. Biol. 223, jeb208975 (2020).

59. Brown-Peterson, N. J. & Heins, D. C. Interspawning interval of wild female three-spined stickleback *Gasterosteus aculeatus* in Alaska. J. Fish Biol. 74, 2299–2312 (2009).

60. Barber, I. Nests as ornaments: revealing construction by male sticklebacks. Behav. Ecol. 12, 390–396 (2001).

61. Government of British Columbia, GeoBC. Hydrological dataset, Geospatial dataset, BC Data Catalogue. Freshwater Atlas - Province of British Columbia https://www2.gov.bc.ca/gov/content/data/geographic-data-services/topographic-data/freshwater (2024).

62. Vermote, E. VIIRS/NPP Vegetation Indices Monthly L3 Global 0.05Deg CMG V002. NASA Land Processes Distributed Active Archive Center 10.5067/VIIRS/VNP13C2.002 (2023).

63. Schönberger, D., & Wang, T. ClimateNAr: R interface to ClimateNA climate data (Version 3.1.0). ClimateNAr R package (2025) doi:10.5281/zenodo.17401570.

64. Schönberger, D. & Wang, T. ClimateNAr R Package (mirror for Reproducible Installation). (Zenodo, 2025). doi:10.5281/ZENODO.17401570.

65. Hollister, J. & Stachelek, J. lakemorpho: Calculating lake morphometry metrics in R. F1000Res. 6, 1718 (2017).

66. Hengl, T., Heuvelink, G. B. M. & Rossiter, D. G. About regression-kriging: From equations to case studies. Comput. Geosci. 33, 1301–1315 (2007).

67. Prudhomme, C. & Reed, D. W. Mapping extreme rainfall in a mountainous region using geostatistical techniques: a case study in Scotland. Int. J. Climatol. 19, 1337–1356 (1999).

68. Omuto, C. T. & Vargas, R. R. Re-tooling of regression kriging in R for improved digital mapping of soil properties. Geosci. J. 19, 157–165 (2015).

69. Manaf, M., Ali, Z. & Scholz, M. Integrating random forest-based regression kriging for analyzing spatial variability of rainfall in arid and semi-arid regions. Sci. Rep. 16, 5298 (2026).

70. Takoutsing, B. & Heuvelink, G. Comparing the prediction performance, uncertainty quantification and extrapolation potential of regression kriging and random forest while accounting for soil measurement errors. Geoderma 428, (2022).

71. Berini, J. L., Runck, B., Vogeler, J., Fox, D. L. & Forester, J. D. Estimates of woody biomass and mixed effects improve isoscape predictions across a northern mixed forest. Front. Ecol. Evol. 11, (2023).

72. Pal, C., Bengtsson-Palme, J., Kristiansson, E. & Larsson, D. G. J. Co-occurrence of resistance genes to antibiotics, biocides and metals reveals novel insights into their co-selection potential. BMC Genomics 16, 964 (2015).

73. Murray, L. M. et al. Co-selection for antibiotic resistance by environmental contaminants. NPJ Antimicrob. Resist. 2, 9 (2024).

74. Larsson, D. G. J. & Flach, C.-F. Antibiotic resistance in the environment. Nat. Rev. Microbiol. 20, 257–269 (2022).

75. Decaestecker, E. et al. Host–parasite ‘Red Queen’ dynamics archived in pond sediment. Nature 450, 870–873 (2007).

76. Nickodem, C. A. et al. Soil management strategies drive divergent impacts on pathogens and environmental resistomes. Sci. Rep.

77. Baquero, F., Tedim, A. P. & Coque, T. M. Antibiotic resistance shaping multi-level population biology of bacteria. Front. Microbiol. 4, 15 (2013).

78. Baquero, F., Martínez, J.-L. & Cantón, R. Antibiotics and antibiotic resistance in water environments. Curr. Opin. Biotechnol. 19, 260–265 (2008).

79. Ng, C. et al. Characterization of metagenomes in urban aquatic compartments reveals high prevalence of clinically relevant antibiotic resistance genes in wastewaters. Front. Microbiol. 8, (2017).

80. Shao, S., Hu, Y., Cheng, J. & Chen, Y. Research progress on distribution, migration, transformation of antibiotics and antibiotic resistance genes (ARGs) in aquatic environment. Crit. Rev. Biotechnol. 38, 1–14 (2018).

81. McCubbin, K. D. et al. Knowledge gaps in the understanding of antimicrobial resistance in Canada. Front. Public Health 9, 726484 (2021).

82. Mustaq, S., Moin, A., Pandit, B., Tiwary, B. K. & Alam, M. *Phyllobacteriaceae*: a family of ecologically and metabolically diverse bacteria with the potential for different applications. Folia Microbiol. (Praha) 69, 17–32 (2024).

83. Puk, K. & Guz, L. Occurrence of *Mycobacterium* spp. in ornamental fish. Ann. Agric. Environ. Med. 27, 535–539 (2020).

84. Gauthier, D. T. & Rhodes, M. W. Mycobacteriosis in fishes: a review. Vet. J. 180, 33–47 (2009).

85. Bolnick, D. I. et al. Individual diet has sex-dependent effects on vertebrate gut microbiota. Nat. Commun. 5, 4500 (2014).

86. Härer, A., Thompson, K. A., Schluter, D. & Rennison, D. J. Associations between gut Microbiota diversity and a host fitness proxy in a naturalistic experiment using threespine stickleback fish. Mol. Ecol. 33, e17571 (2024).

87. Härer, A., Kurstjens, E. & Rennison, D. J. Host traits and environmental variation shape gut microbiota diversity in wild threespine stickleback. *Anim*. Microbiome 7, 67 (2025).

88. Bagamian, K. H., Heins, D. C. & Baker, J. A. Body condition and reproductive capacity of three-spined stickleback infected with the cestode *Schistocephalus solidus*. J. Fish Biol. 64, 1568–1576 (2025).

89. Schwenke, R. A., Lazzaro, B. P. & Wolfner, M. F. Reproduction-immunity trade-offs in insects. Annu. Rev. Entomol. 61, 239–256 (2016).

90. Brotman, R. M., Ravel, J., Bavoil, P. M., Gravitt, P. E. & Ghanem, K. G. Microbiome, sex hormones, and immune responses in the reproductive tract: challenges for vaccine development against sexually transmitted infections. Vaccine 32, 1543–1552 (2014).

91. Braibant, M., Chevalier, J., Chaslus-Dancla, E., Pagès, J.-M. & Cloeckaert, A. Structural and functional study of the phenicol-specific efflux pump FloR belonging to the major facilitator superfamily. Antimicrob. Agents Chemother. 49, 2965–2971 (2005).

92. Aínsa, J. A. et al. Molecular cloning and characterization of Tap, a putative multidrug efflux pump present in *Mycobacterium fortuitum* and *Mycobacterium tuberculosis*. J. Bacteriol. 180, 5836–5843 (1998).

93. Siddiqi, N., et al. *Mycobacterium tuberculosis* isolate with a distinct genomic identity overexpresses a tap-like efflux pump. Infection 32, 109–111 (2004).

94. Tilseth, S., Hansen, T. & Møller, D. Historical development of salmon culture. Aquaculture 98, 1–9 (1991).

95. Morrison, D. B. & Saksida, S. Trends in antimicrobial use in Marine Harvest Canada farmed salmon production in British Columbia (2003-2011). Can. Vet. J. 54, 1160–1163 (2013).

96. Roberts, M. C. & Schwarz, S. Tetracycline and phenicol resistance genes and mechanisms: Importance for agriculture, the environment, and humans. J. Environ. Qual. 45, 576–592 (2016).

97. Shur, K. V. et al. The intrinsic antibiotic resistance to β-lactams, macrolides, and fluoroquinolones of mycobacteria is mediated by the *whiB7* and *tap* genes. Russian Journal of Genetics 53, 1006–1015 (2017).

98. Panda, A., Drancourt, M., Tuller, T. & Pontarotti, P. Genome-wide analysis of horizontally acquired genes in the genus *Mycobacterium*. Sci. Rep. 8, 14817 (2018).

99. Ramón-García, S. et al. Functional and genetic characterization of the Tap efflux pump in *Mycobacterium bovis* BCG. Antimicrob. Agents Chemother. 56, 2074–2083 (2012).

100. Pao, S. S., Paulsen, I. T. & Saier, M. H., Jr. Major facilitator superfamily. Microbiol. Mol. Biol. Rev. 62, 1–34 (1998).

101. Quistgaard, E. M., Löw, C., Guettou, F. & Nordlund, P. Understanding transport by the major facilitator superfamily (MFS): structures pave the way. Nat. Rev. Mol. Cell Biol. 17, 123–132 (2016).

102. Drew, D., North, R. A., Nagarathinam, K. & Tanabe, M. Structures and general transport mechanisms by the major facilitator superfamily (MFS). Chem. Rev. 121, 5289–5335 (2021).

103. Wang, Y., Batra, A., Schulenburg, H. & Dagan, T. Gene sharing among plasmids and chromosomes reveals barriers for antibiotic resistance gene transfer. Philos. Trans. R. Soc. Lond. B Biol. Sci. 377, 20200467 (2022).

104. Carlson, R. E. A trophic state index for lakes1. Limnol. Oceanogr. 22, 361–369 (1977).

105. Goryluk-Salmonowicz, A. & Popowska, M. Factors promoting and limiting antimicrobial resistance in the environment - Existing knowledge gaps. Front. Microbiol. 13, 992268 (2022).

106. Knöppel, A., Näsvall, J. & Andersson, D. I. Evolution of antibiotic resistance without antibiotic exposure. Antimicrob. Agents Chemother. 61, (2017).

107. Clinton, M., Wyness, A. J., Martin, S. A. M., Brierley, A. S. & Ferrier, D. E. K. Sampling the fish gill microbiome: a comparison of tissue biopsies and swabs. BMC Microbiol. 21, 313 (2021).

108. Kozich, J. J., Westcott, S. L., Baxter, N. T., Highlander, S. K. & Schloss, P. D. Development of a dual-index sequencing strategy and curation pipeline for analyzing amplicon sequence data on the MiSeq Illumina sequencing platform. Appl. Environ. Microbiol. 79, 5112–5120 (2013).

109. Quast, C. et al. The SILVA ribosomal RNA gene database project: improved data processing and web-based tools. Nucleic Acids Res. 41, D590–6 (2013).

110. Wood, D. E., Lu, J. & Langmead, B. Improved metagenomic analysis with Kraken 2. Genome Biol. 20, 257 (2019).

111. Camacho, C. et al. BLAST+: architecture and applications. BMC Bioinformatics 10, 421 (2009).

112. Alcock, B. P. et al. CARD 2023: expanded curation, support for machine learning, and resistome prediction at the Comprehensive Antibiotic Resistance Database. Nucleic Acids Res. 51, D690–D699 (2023).

113. Jung, Y.-J., Kim, H.-J. & Hur, M. *Mesorhizobium terrae* sp. nov., a novel species isolated from soil in Jangsu, Korea. Antonie Van Leeuwenhoek 113, 1279–1287 (2020).

114. Fouilloux, C. A. et al. Needle in a haystack: A droplet digital polymerase chain reaction assay to detect rare helminth parasites infecting natural host populations. Molecular Ecology Resources 25, e14131 (2025).

